# Transcriptomic, Specific Marker, and Pathway Analysis of Smooth Muscle Cell Foam Cells Relative to Macrophage Foam Cells in Human Atherosclerosis

**DOI:** 10.64898/2026.05.27.728334

**Authors:** Sima Allahverdian, Yuancheng Mao, Pinhao Xiang, Valentin Blanchard, Aydin Bölük, Patrick A. Hart, Paul Cheng, Daniel Y. Li, Matthew D. Worssam, Uma Thanigai Arasu, Mari Taipale, Miika Kiema, Johanna P. Laakkonen, Tiit Örd, Minna U. Kaikkonen, Clint L. Miller, Thomas Quertermous, Teddy Chan, Gordon A. Francis

## Abstract

**BACKGROUND:** Smooth muscle cells (SMCs) are reported to contribute the majority of cholesterol-overloaded foam cells in human and mouse atheromas. However, the transcriptome, specific markers, and biologic itinerary of SMC foam cells relative to macrophage foam cells have not been determined.

**METHODS:** Transcriptomic analysis by single cell RNA sequencing (scRNA-seq) was performed on fresh coronary segments from heart transplant recipients with early to intermediate stage atherosclerosis. The gene expression pattern of a putative cluster of SMC foam cells was compared to those of cultured SMCs loaded with either aggregated low density lipoprotein (agLDL) or cholesterol bound to methyl-β-cyclodextrin (Chol-MβCD). Candidate markers of SMC foam cells not expressed by macrophage foam cells were validated in ours and publicly available datasets, by spatial transcriptomics and by immunofluorescence microscopy of human atheromas. Pathway analysis was performed using Gene Set Enrichment Analysis Hallmark gene sets.

**RESULTS:** SMC foam cells derived from fibromyocytes were tentatively identified using a panel of markers upregulated with in vitro cholesterol loading of SMCs. agLDL loading reproduced the same transcriptional profile, whereas Chol-MβCD did not reproduce any in vivo SMC state. Top genes highly represented in SMC foam cells included *SERPINE1*, encoding plasminogen activator inhibitor 1 (PAI-1) and *CFH*, complement factor H, which were validated in further human coronary scRNA-seq datasets, by Xenium spatial transcriptomics, and by immunofluorescence microscopy. Relative to macrophage foam cells, SMC foam cells exhibit a distinct biologic itinerary, including activation of extracellular matrix, coagulation and angiogenesis pathways.

**CONCLUSIONS:** SMC foam cells, which are derived from fibromyocytes (“lipomyocytes”), exhibit unique markers and biologic programs that differ markedly from macrophage foam cells in atherosclerotic plaque development. Further understanding of the role of lipomyocytes and their expression of CFH and PAI-1 expression in plaque biology may offer novel therapeutic options to reduce ischemic cardiovascular disease.

**Novelty and Significance:** *What is Known?:*

- Single-cell transcriptomic studies have demonstrated vascular SMCs undergo extensive phenotypic modulation during atherosclerosis, giving rise to fibromyocytes and other intermediate cell states.
- SMC-derived foam cells constitute a major proportion of plaque foam cells.
- Previous studies have shown that SMC-derived foam cells differ from macrophage-derived foam cells in cholesterol handling and lipid metabolism, making their identification using conventional macrophage-associated foam-cell markers challenging, and highlighting the need for SMC foam cell-specific molecular markers.

*What New Information Does This Article Contribute:*

- By integrating published transcriptomic datasets with bulk RNA-sequencing of agLDL-loaded human SMCs, we identify the transcriptional program of human SMC-derived foam cells and demonstrate that these cells are embedded within the fibromyocyte population rather than forming a distinct cell cluster.
- agLDL loading in vitro reproduces the in vivo SMC foam-cell phenotype, whereas loading with cholesterol bound to cyclodextrin induces an inflammatory state not found in any in vivo SMC state in human atherosclerosis.
- *CFH* and *SERPINE1* were identified as specific markers of SMC-derived foam cells and validated by immunostaining and Xenium spatial transcriptomics, providing robust markers for identification of these cells in human atherosclerotic plaques.
- The top transcriptional programs activated in SMC foam cells differ completely from macrophage foam cells and include extracellular matrix remodeling, complement regulation, coagulation and angiogenesis. In this study, we mapped smooth muscle foam cells to the fibromyocyte cluster of arterial SMC subtypes in human coronary artery atheromas. In vitro loading of smooth muscle cells with aggregated low-density lipoprotein, the presumed physiologic substrate for foam cell formation during plaque development, induced a gene expression pattern that localized to the same fibromyocyte cluster. The proposed sequence of SMC foam cell formation is that fibromyocytes present in human pre-atherosclerotic intima generate the proteoglycans that bind lipoproteins and allow their uptake by fibromyocytes to generate cholesterol-loaded “lipomyocytes”. Lipid loading of fibromyocytes drives expression of specific markers and transcriptional programs, but with an overall gene expression pattern that continues to resemble fibromyocytes. We demonstrate here that lipomyocytes express the specific markers *CFH* and *SERPINE1*, and exhibit activation of a completely different set of pathways relative to macrophage foam cells in the plaque. These findings provide evidence for the unique phenotype and specific markers of lipomyocytes relative to macrophage foam cells in human atherosclerosis, which will be critical in evaluating the response of these different types of foam cell to novel therapies.

## INTRODUCTION

Cholesterol accumulation in intimal foam cells is the defining biochemical hallmark of atherosclerosis, and the principal therapeutic target for preventing and regressing the atheromas that lead to ischemic vascular events.^1^ While plaque cholesterol accumulation has in recent years been attributed largely to monocyte-derived macrophages,^2,3^ literature starting from the time of Virchow in the 1850’s and in the 1990’s suggested arterial intimal smooth muscle cells (SMCs) are the main cell type accumulating excess lipid in early atherogenesis.^4,5^ Initial accumulation of cholesterol in the human artery wall is mediated by retention of apolipoprotein B-containing lipoproteins through their binding to extracellular matrix proteoglycans secreted by SMCs that populate the intima in the preatherosclerotic diffuse intimal thickening stage.^6^ In human atherogenesis, subsequent lipoprotein uptake likely occurs first in SMCs and then later in macrophages, leading to foam cell formation.^6^ In contrast, studies in mice--which lack a preatherosclerotic intima and require a highly inflammatory, hyperlipidemic state initially driven by monocytes and macrophages to develop atherosclerosis--have been interpreted as indicating that macrophages contribute the bulk of foam cells within the plaque.^2,3^

Studies in recent years from our group and others of human^7^ and mouse^8–10^ atheromas suggested SMCs, rather than macrophages, contribute the majority of foam cells in the plaque. While several publications have outlined the biologic itinerary and attempted to define the transcriptome of macrophage foam cells,^2,11,12^ the gene expression pattern of SMC foam cells and their biologic role in the plaque have not been determined. When compared to macrophage foam cells, SMC foam cells exhibit major differences their ability to metabolize excess cholesterol, including reduced expression of the rate-limiting cholesterol efflux promoter ATP-binding cassette transporter A1 (*ABCA1*),^7,8,13^ as well as regulators of ABCA1 expression, sterol-27 hydroxylase (*CYP27A1*) and liver X receptor alpha (*LXRα*).^14^ The sole lysosomal enzyme mediating hydrolysis of lipoprotein- and cell-derived cholesteryl esters (CE), lysosomal acid lipase, encoded by *LIPA*, is expressed at very low levels in arterial SMCs and SMC foam cells when compared to human macrophages and macrophage foam cells.^14^ As a result, SMC foam cells retain much of the ingested CE present on lipoproteins within lysosomes, rather than hydrolyzing the CE and delivering unesterified cholesterol to the endoplasmic reticulum to be reformed for storage in the cytoplasm as CE droplets, as seen in macrophages.^14,15^ SMC foam cells therefore lack key machinery to metabolize and efflux ingested CE and free cholesterol. This suggests they represent a potentially regression-resistant repository of cholesterol in the artery wall, even after marked reduction of plasma low density (LDL) and other atherogenic lipoproteins with statins, ezetimibe and PCSK9 inhibitors.

While macrophage foam cells are widely believed to promote plaque inflammation, instability, necrosis and increased susceptibility to rupture,^11,16,17^ the contribution of SMC foam cells to these and other aspects of plaque biology remain largely unknown. Single-cell RNA sequencing has been employed to investigate the heterogeneity of both macrophages and SMCs in human and mouse models of atherosclerosis. Mosquera *et al.* performed a meta-analysis of several scRNAseq datasets derived from human coronary and carotid atheromas, and used a panel of lipid-metabolism-associated genes to tentatively identify a SMC cluster they designated “foam-like”.^18^ Independently, Bashore *et al.*, using a human carotid artery scRNAseq dataset, also used a panel of lipid metabolism genes to identify an SMC cluster they reported as foam cells, expressing the SMC differentiation marker smoothelin and the GPI-anchored cell-surface glycoprotein CD90, encoded by *THY1*.^19^ A recent lineage-tracing mouse study integrating multimodal single-cell analyses identified heterogeneous populations of modulated VSMCs, including a lipid uptake–associated subpopulation, and implicated BHLHE40 as a transcriptional regulator of VSMC phenotypic modulation and foam cell formation.^10^ In the current study, we utilized a set of genes upregulated in in vitro studies of SMC foam cell development and combined this with single-cell RNA sequencing to identify a putative SMC foam cell cluster in human coronary arteries with early to intermediate stage atherosclerosis from patients undergoing heart transplant. We further utilized bulk RNA sequencing of cultured human aortic SMCs loaded with either aggregated LDL (agLDL) or cholesterol bound to methyl-β-cyclodextrin (Chol-MβCD) and mapped the top genes upregulated in these cells onto our scRNAseq of coronary arteries. The results presented here demonstrate that SMC-derived foam cells are embedded within the fibromyocyte population rather than forming a distinct transcriptional cluster. Because these lipid-laden fibromyocytes exhibit a unique molecular phenotype, including expression of *CFH* and *SERPINE1*, we propose the term “lipomyocyte” to describe this specialized SMC foam-cell state. Pathway analysis of SMC foam cells is also performed and compared to macrophage foam cells. Our findings indicate that lipomyocytes in early-to intermediate-stage atherosclerosis are not “macrophage-like”, but instead constitute a unique repository of excess cholesterol in the plaque with their own specific markers and biologic itinerary. Given that SMCs appear to constitute the majority of foam cells in atheromas, understanding their biological trajectory is crucial to determining whether they represent a unique therapeutic target to reduce major ischemic cardiovascular events.

## METHODS

Detailed Methods are available in the Supplemental Materials.

### Data Availability

Single-cell RNA sequencing data will be deposited in the National Center for Biotechnology Information Gene Expression Omnibus (http://www/ncbi.nlm.nih.gov/geo).

### Human coronary artery cell dissociation

Fresh human coronary arteries were dissected from explanted hearts of transplant recipients obtained from the Bruce McManus Cardiovascular Biobank with consent from patients and approval from the University of British Columbia Research Ethics Board. The proximal to mid-right coronary artery (RCA) was dissected and cleaned to remove the perivascular adipose tissue and adventitia. After several washes with cold PBS, the lumen was scraped to remove endothelial cells, and the artery was cut into small pieces. Minced tissue was digested for 60 minutes at 37°C with a cocktail of collagenase (0.3%) and elastase (0.1%) from Sigma Aldrich.^20^ The digested coronary artery was passed through a 40μm cell strainer and spun down at 600xg for 5 minutes to collect the cell pellet. The enzyme solution was then discarded, and the cells were resuspended in fresh DMEM. Cell viability was tested by staining with Trypan Blue. Samples were then transferred on ice to the Sequencing Core at the UBC Biomedical Research Centre for single-cell RNA-sequencing.

### Single-Cell RNA Sequencing and Data Analysis

Single-cell RNA sequencing libraries were generated using the 10x Genomics Chromium platform and processed with Cell Ranger. Downstream quality control, normalization, clustering, and cell-type annotation were performed in R using <u>Seurat</u> (version 4.3.0). Low-quality cells were excluded based on the number of detected genes, UMI counts, and mitochondrial transcript content. Data were normalized, highly variable genes were identified, and cells were analyzed using principal component analysis, shared nearest-neighbor clustering, and UMAP visualization. Cluster-specific marker genes were identified using FindAllMarkers, and differential gene expression between selected clusters was performed using the MAST statistical framework. Additional details of sample processing, quality control criteria, and computational analysis are provided in the Supplemental Methods.

### Integration of Samples with Integrative Atlas of Atherosclerotic Vascular Tissue

scRNA-seq libraries generated using early to intermediate coronary artery lesions were processed using the 10x Genomics CellRanger Count function. Aligned libraries were processed using the scRNAUtils for removal of low-quality cells, doublets, and ambient RNA.^18^ Cells with median absolute deviation (MAD) > 1 for mitochondrial, ribosomal, and hemoglobin content were removed. Data was normalized and Harmony was used for data integration.^21^ Major Leiden clusters identified in UMAP embeddings were annotated based on expression of canonical marker features (*C1QA*: Macrophage, *MYH11*: VSMC, *VWF*: Endothelial, *FBLN1*: Fibroblast, *IL32*, *KIT:* Mast Cells). The data was then subset for VSMC subcluster annotation. A reference dataset, MetaPlaq v2 (Unpublished), was used for label transfer of VSMC subpopulation annotations. The reference dataset was filtered for VSMCs, representing 26,771 cells across contractile, fibromyocyte, fibrochondrocyte, and foam VSMC annotations. MetaPlaq v2 was filtered for libraries containing greater than 200 VSMCs prior to label transfer and the query libraries were concatenated with the reference. The dataset was subset to the 3000 most highly variable genes in Scanpy and the raw counts data was used for subsequent integration.

The scArches workflow was applied for label transfer from the reference to unlabeled VSMC cells in the query dataset.^22^ For the scArches workflow SCVI model, 2 layers and the dimensionality of latent space was set to n_latent=30; default parameters were used otherwise. After integration with the reference, Wilcoxon Rank Sum Test was applied to identify marker features for each VSMC subcluster. After label transfer, VSMC subcluster labels were used to annotate all VSMCs in the Harmony integrated dataset containing 5 broad cell annotations (VSMC, macrophage, endothelial, fibroblast, mast cells).

### Foam Cell Module Gene Scoring using MetaPlaq v2

Gene sets derived from bulk RNA-sequencing of ag-LDL treated VSMCs, Chol-MβCD treated VSMCs, and a set curated from the literature were used to score profiles of individual VSMCs in MetaPlaq v2. Gene scores were calculated in SCANPY.^23^ Visual inspection of scores was performed to assess their enrichment in disease-relevant VSMC subtypes. After gene scoring, a Wilcoxon rank sum test was used to identify marker features for each individual VSMC subset. Genes significantly upregulated in foam VSMCs were then intersected with the gene sets to prioritize candidate regulators and marker features of foam VSMCs.

### Bulk RNA-seq Data Acquisition, Quality Control, and Processing

Total RNA extracted from agLDL- and Chol-MβCD-loaded and control SMCs was submitted to the Sequencing Core at the UBC Biomedical Research Centre for library preparation and sequencing. Sequencing reads were aligned to the human reference genome, and gene-level counts were generated for downstream analysis. Differential gene expression and principal component analysis were performed in R using <u>DESeq2</u> after variance-stabilizing transformation. Genes with an adjusted p-value < 0.05 were considered significantly differentially expressed. Additional details of library preparation, read processing, and statistical analysis are provided in the Supplemental Methods.

### Comparison of bulk RNA-seq profiles with single-cell SMC states

To determine the transcriptional identity of cultured human SMCs prior to cholesterol loading, bulk RNA-seq data from BSA-treated control SMCs were compared with pseudobulk gene expression profiles generated from human coronary artery SMC clusters identified by single-cell RNA sequencing. For each SMC cluster, normalized gene expression values were averaged across all cells to generate pseudobulk expression profiles. The top 2,000 highly variable genes were identified from the single-cell dataset using the FindVariableFeatures function in Seurat (selection method = “vst”), excluding mitochondrial and ribosomal genes. Genes present in both the bulk RNA-seq and pseudobulk datasets were retained for analysis. Spearman correlation coefficients were then calculated between the bulk RNA-seq profile of BSA-treated SMCs and each SMC cluster pseudobulk profile. Correlation coefficients were used to assess transcriptomic similarity between cultured SMCs and the in vivo SMC states.

### Xenium Spatial Transcriptomic Processing and Analysis

Human coronary tissue samples (n = 4) were obtained from patients with cardiomyopathy undergoing heart transplantation at the Stanford University Human Biorepository Tissue Research Bank under approval from the Stanford Institutional Review Board (Protocol No. 32769). All procedures involving human tissue were conducted in accordance with the Declaration of Helsinki and applicable institutional and national guidelines.

Coronary artery samples were collected at the time of orthotopic heart transplantation and fixed in formalin for 12–24 hours at 4°C before paraffin embedding. Xenium spatial transcriptomic analysis was performed on formalin-fixed, paraffin-embedded sections from the left anterior descending or right coronary arteries. Tissue sectioning, placement on Xenium slides, and sample processing were performed according to the manufacturer’s recommended protocol (10x Genomics CG000760, Rev A). The probe panel included 5,001 pre-designed pan-tissue genes and 100 custom add-on genes selected based on coronary artery disease single-cell RNA-seq studies.^24^ Samples were imaged and processed using Xenium Analyzer with instrument software version 4.0.1.4 and onboard analysis version 3.3.1.1. Multimodal cell segmentation was performed using nuclear, boundary, and interior staining.

### Quality Control, Integration, and Cell Type Identification

Data were analyzed in R using Seurat version 5.2.1. Transcript molecules with Phred-scaled quality values <20 were excluded before generation of cell-by-gene count matrices. Genes expressed in fewer than 10 cells were removed. Cells were excluded if they contained fewer than 20 transcripts or had segmented areas smaller than 10 µm² or larger than 200 µm².

Following quality control, data were log-normalized and scaled. Data integration across samples was performed using Harmony,^25^ and clustering was conducted using the Leiden algorithm.^26^

### Gene Set–Based Module Scoring

Gene set–based module scores were calculated using Seurat’s AddModuleScore function to quantify expression of predefined fibromyocyte, macrophage, and foam macrophage signatures. Cells were classified as fibromyocytes when the fibromyocyte score was >0.1 and both macrophage-related scores were <0. Cells were classified as foamy macrophages when both macrophage and foam macrophage scores were >0.1 and the fibromyocyte score was <0. Cells were classified as macrophages when the macrophage score was >0.1, the foam macrophage score was <0, and the fibromyocyte score was <0. Cells that did not meet these criteria were left unassigned.

### Pathway Enrichment Analysis

Pathway enrichment analyses were performed using two complementary approaches. Gene Ontology (GO) Biological Process enrichment analysis was used to identify pathways significantly enriched among selected differentially expressed genes.^27,28^ Gene set enrichment analysis (GSEA) was performed on ranked gene lists to identify pathways enriched across the full spectrum of gene expression changes without requiring an arbitrary significance cutoff.^29^

For GO enrichment analysis, significantly upregulated genes were analyzed using Gene Ontology (Biological Process) annotations, and pathways were ranked according to false discovery rate (FDR)-adjusted significance. For GSEA, genes were ranked by average log2 fold change and analyzed using the fgsea package in R with Hallmark gene sets from the Molecular Signatures Database (MSigDB). Pathways with an FDR < 0.05 were considered significantly enriched.

Leading-edge genes were examined to identify genes contributing most strongly to the enrichment of individual pathways.

### Statistics

All statistical analyses were performed in R (version 4.2.3). Differential gene expression in single-cell RNA-seq data was assessed using the MAST framework implemented in Seurat, with Bonferroni-adjusted p-values. Bulk RNA-seq differential expression was performed using DESeq2 with Benjamini–Hochberg false discovery rate (FDR) correction. Pathway enrichment analyses were considered significant at an FDR-adjusted p-value < 0.05. Bulk RNA-seq experiments included four independent biological replicates per treatment group. Sample sizes and statistical tests are indicated in the corresponding figure legends where applicable. Unless otherwise stated, adjusted p-values < 0.05 were considered statistically significant.

## RESULTS

### Single-cell RNA Sequencing of Diverse Vascular Cell Populations and Smooth Muscle Cell Phenotypic States in Human Coronary Atherosclerotic Plaques

To characterize smooth muscle cell foam cells among the cells comprising early-to intermediate-stage human coronary atherosclerosis, noncalcified right coronary artery plaque segments were obtained from patients undergoing heart transplantation for nonischemic cardiomyopathy (clinical characteristics are provided in Supplemental Table 1). Tissue was processed within 2 hours of explant, enzymatically dissociated into single-cell suspensions, and subjected to single-cell RNA sequencing using 10x Genomics Chromium Single Cell 3ʹ Reagent Kit v3. After quality control, 4,812 cells from three donors were retained for analysis in R using Seurat (version 4.2.3). Unsupervised clustering identified 15 transcriptionally distinct cell populations, including six smooth muscle cell (SMC) clusters, as well as pericytes, fibroblasts, macrophages, endothelial cells, mast cells, T cells, and NK cells (Figure 1A).

**Figure 1.**
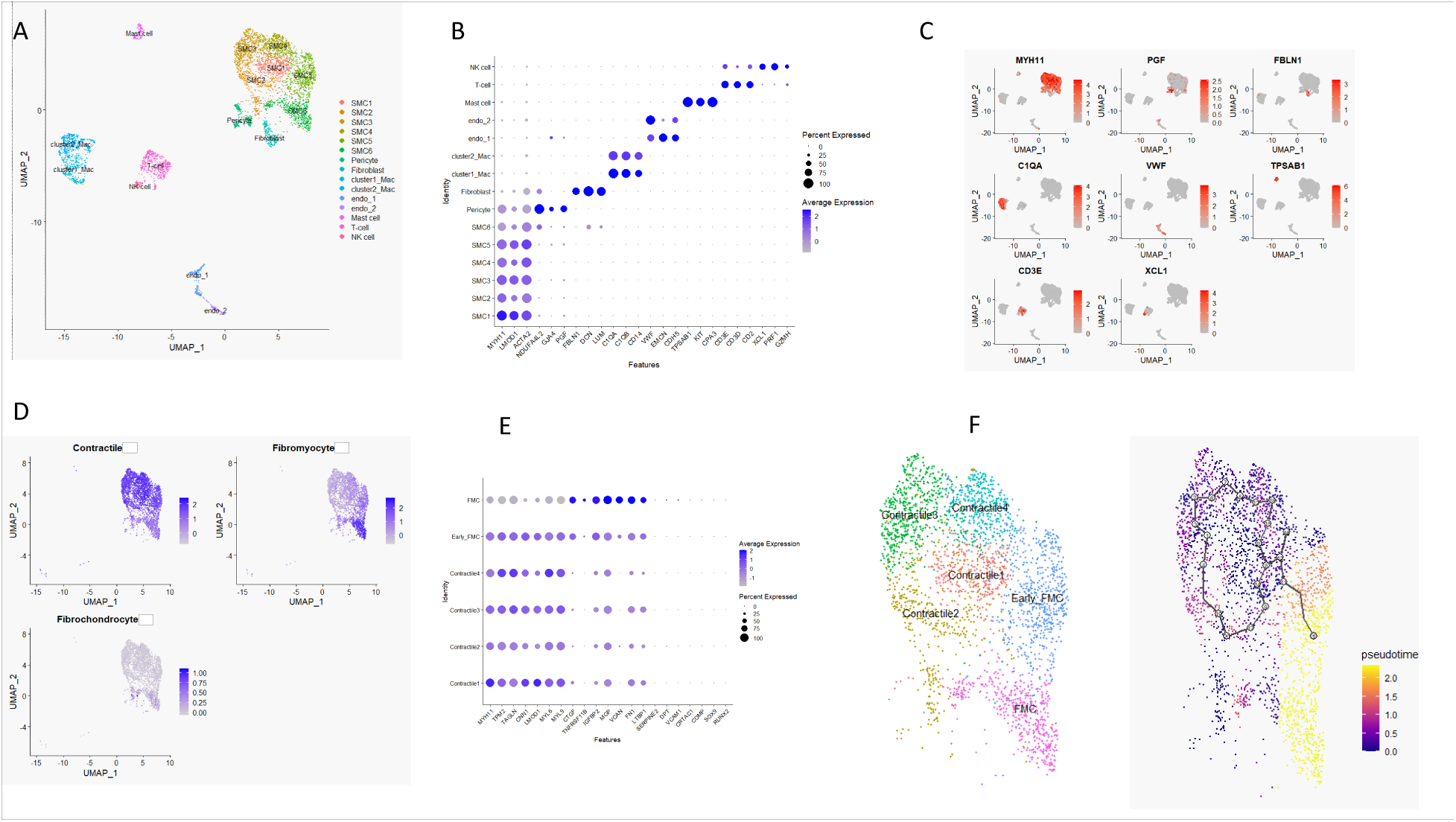
Hierarchical annotation and phenotypic continuum of vascular cell populations with smooth muscle cell (SMC) state transitions. (**A**) UMAP projection of single-cell RNA sequencing (scRNA-seq) data from human coronary artery samples (*n* = 3), showing major vascular and immune cell populations identified by unsupervised clustering using Seurat. **(B)** Dot plot of canonical lineage markers used for Level-1 cell type annotation. Dot size represents the percentage of cells expressing each gene, and color indicates average scaled expression. **(C)** Feature plots of representative marker genes validating cell type identities across clusters, including smooth muscle cells, fibroblasts, endothelial cells, immune cells, and other minor populations. **(D)** UMAP projections displaying module scores for previously described SMC transcriptional programs, highlighting enrichment of contractile, transitional, fibromyocyte, and fibrochondrocyte gene signatures across the SMC compartment. **(E)** Dot plot summarizing Level-2 annotation of SMC subclusters based on marker gene expression and transcriptional programs. SMC1–SMC4 are annotated as Contractile1– Contractile4, SMC5 and SMC7 as Transitional1–Transitional2, and SMC6 as Fibromyocyte; fibroblasts are included for reference. **(F)** UMAP visualization of SMC Level-2 states with inferred pseudotime trajectories overlaid, illustrating a continuous spectrum of SMC phenotypic plasticity from contractile states through transitional intermediates toward fibromyocyte and fibroblast-like states.

Cell identities were assigned using canonical lineage markers visualized by dot plots and feature plots (Figure 1B–C). SMCs expressed *MYH11* and *ACTA2*, pericytes expressed *GJA4* and *PGF*, fibroblasts *FBLN1*, endothelial cells *VWF*, macrophages *C1QA*, mast cells *TPSAB1*, T cells *CD3E*, and NK cells *XCL1*. Marker genes for each cluster were identified using Seurat FindAllMarkers and are provided in Supplemental Table 2. Major cell populations were detected in broadly similar proportions across all three samples (Supplemental Figure 1A).

To characterize SMC heterogeneity, module scores were calculated using previously defined contractile, fibromyocyte, and fibrochondrocyte gene signatures from the MetaPlaq atlas.^18^ Distinct enrichment patterns across SMC clusters revealed a continuum of phenotypic states rather than discrete cell types (Figure 1D). Based on marker expression and module scores, SMC1–SMC4 were classified as Contractile1–Contractile4, SMC5 as Early Fibromyocytes, and SMC6 as Fibromyocytes (Figure 1E). Fibrochondrocyte signatures were present but did not define a distinct cluster, consistent with the noncalcified nature of the lesions. Gene Ontology Biological Process analysis of cluster markers identified distinct functional programs across the SMC continuum (Supplemental Table 3). Contractile states were enriched for muscle contraction, cytoskeletal organization, and energy metabolism, whereas early fibromyocytes were associated with migration, wound healing, and growth factor responses. Fibromyocytes were strongly enriched for extracellular matrix organization, cell adhesion, and transforming growth factor-β signaling, consistent with a matrix-remodeling phenotype.

To validate these annotations coming from atheromas of hearts with nonischemic cardiomyopathy, SMCs were mapped to the MetaPlaq v2 vascular reference atlas using scANVI, demonstrating strong concordance with reference-defined contractile and fibromyocyte states (Supplemental Figure 1B&1C) derived primarily from carotid and coronary samples of patients with atherosclerotic vascular disease. Finally, pseudotime analysis using Monocle 3 revealed a continuous progression from contractile SMCs through transitional intermediates toward fibromyocyte and fibroblast-like phenotypes (Figure 1F).

### Literature-guided Transcriptomic Mapping of SMC Foam Cells

To determine whether a SMC population in our dataset exhibits a transcriptional program consistent with foam cells, we examined the expression of previously reported SMC foam cell– associated gene signatures derived from experimental and human atherosclerosis studies. We first curated a gene set from prior studies describing markers associated with SMC lipid accumulation and foam cell formation following loading with oxidized LDL, including extracellular matrix (ECM)-related genes linked to SMC-derived foam cells,^30^ as well as LDL receptor–related protein 1 (*LRP1*), a mediator of modified lipoprotein uptake in SMCs.^31^ Module scoring of this gene set revealed the strongest enrichment within the fibromyocyte cluster, with additional enrichment observed in fibroblasts (Figure 2A).

**Figure 2.**
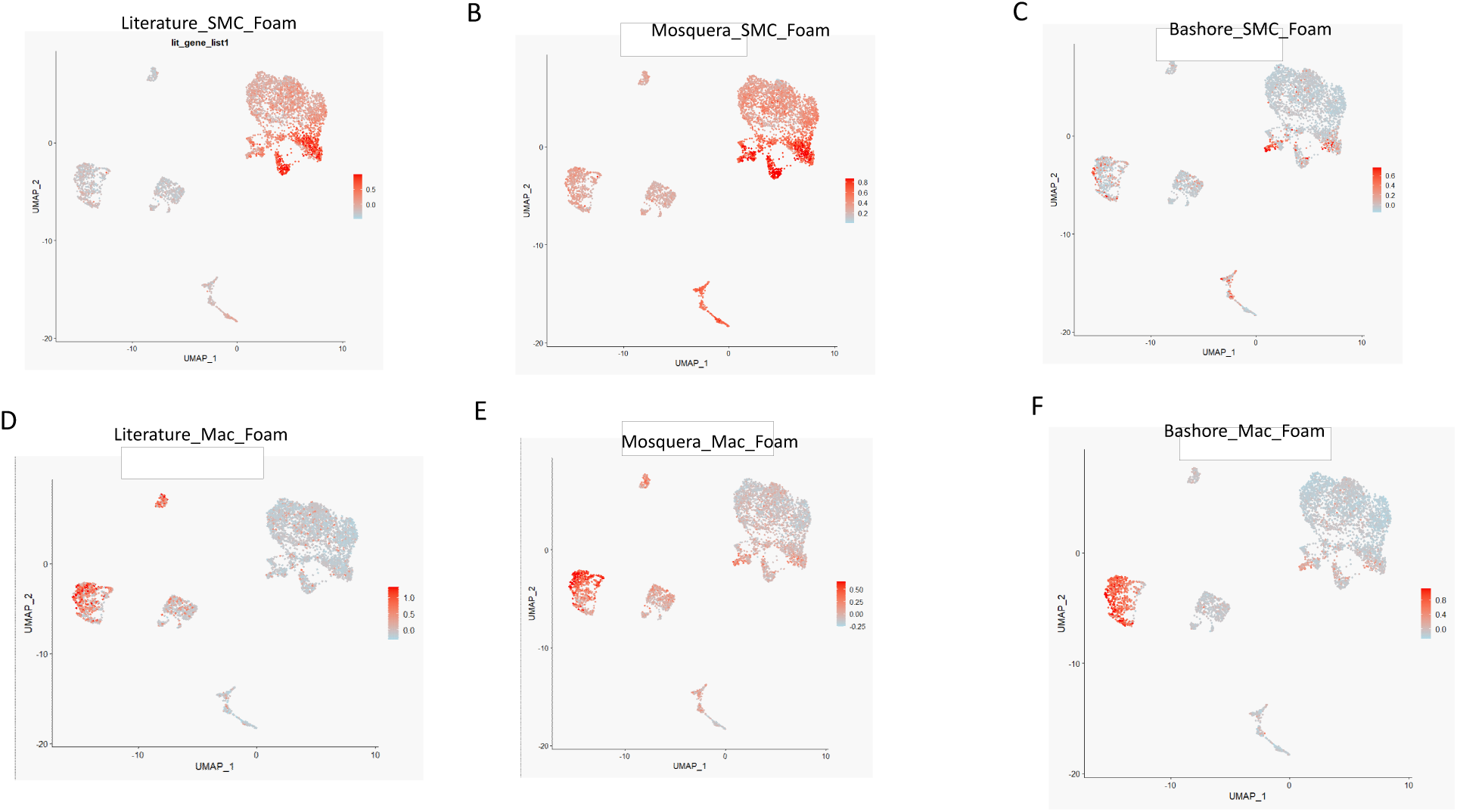
Mapping of foam cell–associated gene signatures onto single-cell transcriptomic data. Module scores were calculated using AddModuleScore and visualized on UMAP embeddings. **(A–C)** SMC foam cell–associated gene signatures. **(A)** Literature-curated SMC gene set including ECM-related genes and LRP1. **(B)** SMC foam cell–like gene signature from Mosquera et al.^18^ **(C)** SMC foam cell–associated gene signature from Bashore et al.^19^ **(D–F)** Macrophage foam cell–associated gene signatures. **(D)** Literature-derived macrophage gene set. **(E)** Macrophage gene signatures from Mosquera et al.^18^ **(F)** Macrophage gene signature from Bashore et al.^19^ Color scale indicates relative module score.

We next examined a SMC foam cell–like gene signature reported by Mosquera *et al.* in their integrative single-cell meta-analysis of human atherosclerosis.^18^ Mapping of this gene set onto our scRNAseq data demonstrated similar enrichment predominantly within fibromyocyte-like SMCs, with additional enrichment observed in pericytes and fibroblasts (Figure 2B), indicating partial overlap between foam cell–associated transcriptional programs and intermediate SMC states undergoing phenotypic modulation. To further validate these observations, we analyzed a third foam cell–associated gene signature derived from the SMC foam cell–like population described by Bashore *et al.* in a high-dimensional multimodal study of human carotid atherosclerosis.^19^ This more limited gene set showed a similar pattern of enrichment, again localizing primarily to fibromyocyte-like SMCs, with lower enrichment in pericytes and fibroblasts (Figure 2C).

Notably, although fibroblasts exhibited a foam cell–associated transcriptional signature, this likely reflects shared matrix remodeling and stress-response programs rather than true lipid accumulation. Immunofluorescence analysis of human coronary atheromas demonstrated the majority of Fibulin-1 (FBLN1)-positive fibroblasts in the adventitial layer, very few in the intima, and absence of BODIPY-positive lipid accumulation in FBLN1-positive fibroblasts in the intimal layer (Supplemental Figure 2A–C). This indicates the specificity of non-macrophage foam cells to the fibromyocyte cluster of SMCs.

In parallel, we evaluated macrophage foam cell–associated transcriptional programs using gene signatures derived from prior literature^9,14,32^ as well as the independent single-cell studies of Mosquera *et al.* and Bashore *et al*.^18,19^ The full list of genes contributing to these analyses is provided in Table 1. Module scoring of a literature-curated macrophage foam cell gene set demonstrated strong enrichment within macrophage populations (Figure 2D).

**Table 1.**
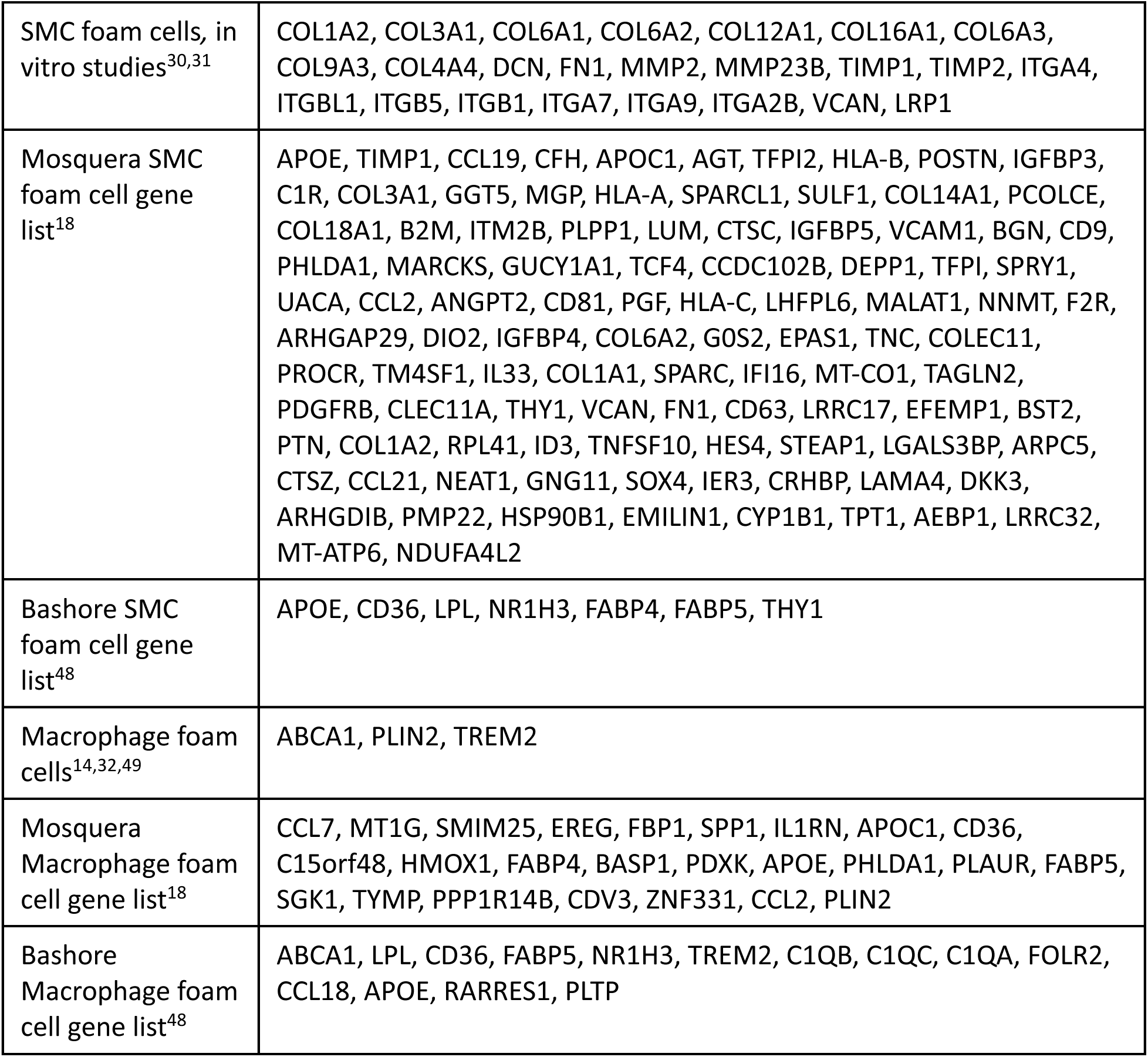

Macrophage foam cell gene signatures derived from Mosquera^18^ and Bashore^19^ showed a similar pattern, with enrichment localized to both macrophage clusters (Figure 2E–F). For the Mosquera-derived signature, we refined the gene set by selecting genes with log2FC > 0.8 and pct.2 < 0.25 and excluding pan-macrophage markers to better capture foam cell–specific programs. Using this refined set, enrichment was more pronounced in cluster2_Mac, supporting its identification as the macrophage subset most closely associated with a foam cell–like phenotype. In contrast, the Bashore-derived gene set showed enrichment across both macrophage clusters, suggesting that both populations exhibit foam cell–associated features in this dataset. Macrophage-derived gene signatures showed minimal enrichment within SMC populations, and vice versa. Together, these analyses consistently indicate that fibromyocytes represent the population most enriched for SMC foam cell–associated transcriptional programs, and that these differ markedly from macrophage foam cells.

### Transcriptomic Profiling of Aggregated-LDL-loaded SMCs Identifies Gene Signatures of SMC-derived Foam Cells

Cholesterol loading using cholesterol–methyl-β-cyclodextrin (Chol-MβCD) has been a widely used method to induce lipid accumulation in SMCs for gene expression and functional analysis in vitro.^33–35^ To investigate further the transcriptional programs associated with SMC-derived foam cells, human vascular SMCs were treated with aggregated LDL (agLDL, 100 μg/mL), considered to be the most likely physiologic substrate for foam cell formation in the intima,^36^ or Chol-MβCD (10 μg/mL) for 72 hours and subjected to bulk RNA sequencing alongside BSA-treated SMC controls (n = 4 per group). Principal component analysis (PCA) demonstrated clear transcriptional separation between the two treated and control SMC groups (Figure 3A). To determine the phenotypic identity of cultured SMCs prior to cholesterol loading, we compared the transcriptomic profiles of BSA-treated control cells with the human coronary artery SMC clusters identified above. BSA-treated SMCs showed the greatest transcriptomic similarity to the fibromyocyte cluster (Supplemental Figure 3), indicating that cultured SMCs already exhibit a fibromyocyte-like transcriptional state before lipid loading.

**Figure 3.**
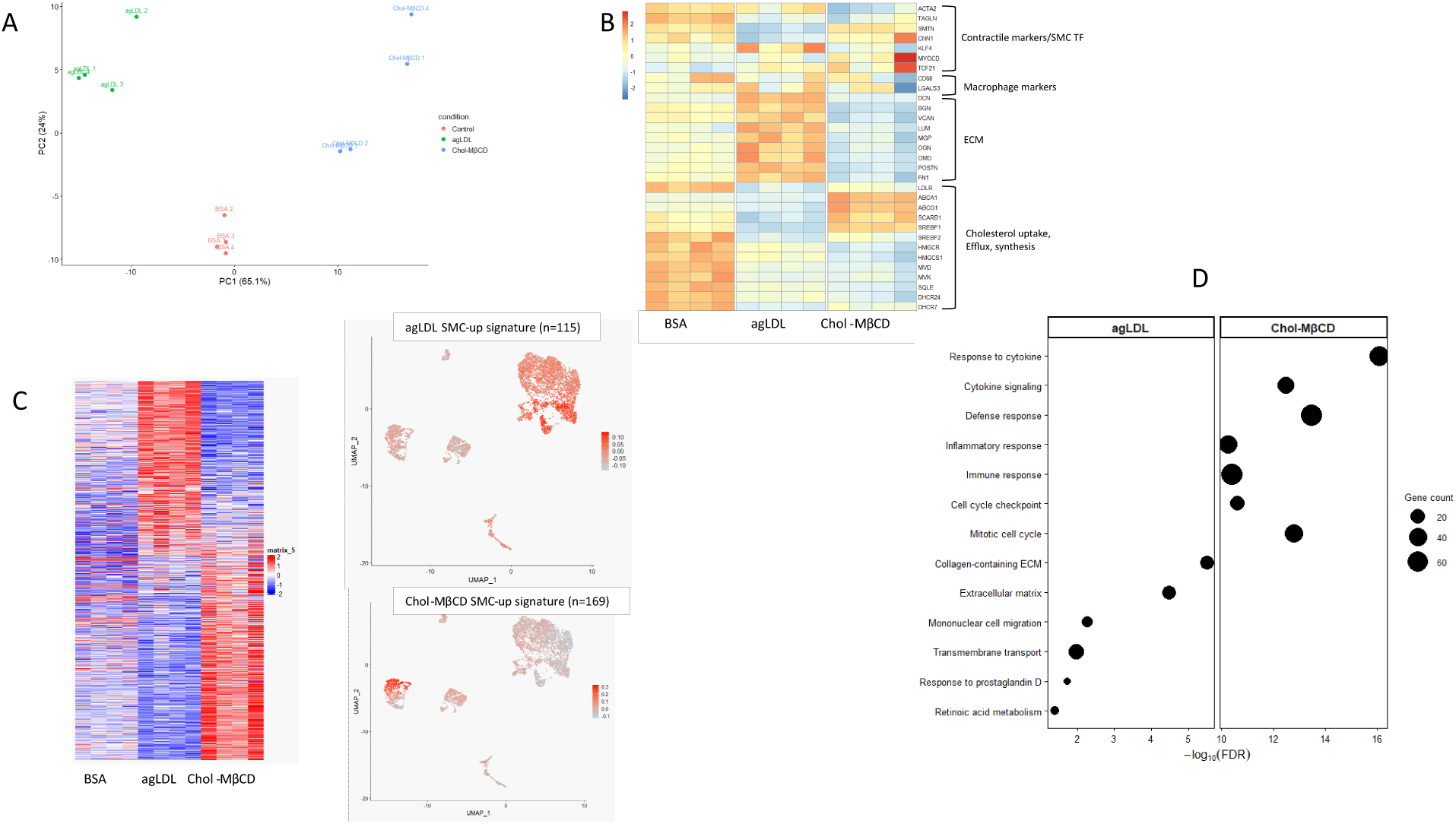
Transcriptomic profiling and pathway analysis of agLDL- and Chol-MβCD-loaded SMCs. Human vascular SMCs were treated with 100 μg/mL agLDL or 10 μg/mL Chol-MβCD for 72 h (n = 4 per condition). **(A)** Principal component analysis (PCA) demonstrates clear separation between treated and control SMCs, indicating distinct transcriptional responses to agLDL and Chol-MβCD. **(B)** Heatmap of representative gene sets across treatment conditions. Genes are organized into functional categories, including contractile SMC markers and transcription factors, macrophage-associated genes, extracellular matrix (ECM) components, and genes involved in cholesterol uptake, efflux, and biosynthesis. **(C)** Left: Heatmap of differentially expressed genes (log₂FC > 1, adjusted P value < 0.05) showing global transcriptional changes induced by agLDL and Chol-MβCD treatment. Right: Module scores derived from agLDL-induced (n = 115 genes) and Chol-MβCD–induced (n = 169 genes) SMC gene signatures projected onto a human coronary artery single-cell RNA-seq dataset. **(D)** Gene Ontology (GO: Biological Process) enrichment analysis of upregulated genes in agLDL- and Chol-MβCD-treated SMCs. Dot size represents the number of genes contributing to each pathway, and the x-axis indicates enrichment significance (−log₁₀ FDR).

To examine specific transcriptional programs associated with cholesterol loading, we evaluated the expression of representative genes related to SMC phenotype and lipid handling (Figure 3B). Cholesterol loading altered the expression of several genes associated with SMC phenotype. In agLDL-treated SMCs, expression of the canonical contractile markers *TAGLN*, *MYH11*, and *CNN1* was reduced, whereas *ACTA2* expression remained largely unchanged. In contrast, Chol-MβCD treatment significantly reduced *ACTA2* and *TAGLN* expression, while *SMTN* and *CNN1* expression were not significantly affected. Although cultured SMCs already exhibited a fibromyocyte-like transcriptional profile prior to lipid loading (Supplemental Figure 3), agLDL treatment further increased expression of the transcription factor KLF4, a key regulator of SMC phenotypic plasticity and phenotypic switching^37^, whereas Chol-MβCD treatment had no significant effect. Expression of transcriptional regulators associated with SMC lineage control also differed between the two treatments. The transcriptional co-activator *MYOCD* (myocardin), a central regulator of the contractile SMC gene program,^38^ and *TCF21*, a transcription factor implicated in SMC phenotypic switching toward fibroblast-like states during atherosclerosis,^24^ showed no significant change in agLDL-treated cells but were upregulated following Chol-MβCD treatment. Expression of macrophage-associated markers *CD68* and *LGALS3* did not change significantly in either treatment condition.

While agLDL treatment significantly increased the expression of extracellular matrix remodeling genes, Chol-MβCD treatment was associated with downregulation of these genes, indicating highly distinct transcriptional responses to these two cholesterol-loading methods. Expression of genes involved in cholesterol metabolism showed both shared and distinct responses between treatments. *LDLR* expression was reduced in both agLDL- and Chol-MβCD– treated SMCs. Similarly, genes involved in cholesterol biosynthesis were broadly downregulated under both conditions. In contrast, the expression of cholesterol efflux transporters *ABCA1* and *ABCG1* showed divergent regulation, consistent with differential activation of cholesterol handling pathways in response to agLDL versus Chol-MβCD.

Differential expression analysis using DESeq2 identified genes significantly up- or down-regulated (adjusted P < 0.05) in cholesterol-loaded SMCs compared with control cells (complete gene lists are provided in Supplemental Table 4 and 5). These differentially expressed genes revealed broad transcriptional reprogramming that differed markedly between agLDL and Chol-MβCD treatments (Figure 3C, left). To determine whether these lipid-induced transcriptional programs correspond to SMC-derived foam cell populations in human atherosclerotic plaques, we generated gene signatures from the significantly upregulated genes in each treatment group (log₂ fold change > 1 and adjusted P < 0.05; agLDL signature, n = 115 genes; Chol-MβCD signature, n = 169 genes) and projected them onto our human coronary artery single-cell scRNA-seq dataset using AddModuleScore. Module scoring revealed that the agLDL-induced SMC signature was enriched primarily in fibromyocyte and fibroblast clusters, similar to the approach shown in Figure 2, whereas the Chol-MβCD-derived signature showed strongest enrichment in macrophage rather than SMC clusters (Figure 3C, right).

Gene Ontology (GO: Biological Process) enrichment analysis of the upregulated genes further revealed distinct pathway activation between agLDL- and Chol-MβCD–treated SMCs (Figure 3D). agLDL treatment was associated with enrichment of pathways related to extracellular matrix organization, collagen-containing extracellular matrix, and mononuclear cell migration, whereas Chol-MβCD treatment enriched pathways involved in cytokine-mediated signaling, inflammatory and immune responses, and cell cycle–related processes. Complete pathway enrichment results are provided in Supplemental Table 6 and 7. Together, these findings indicate that cultured human SMCs already exhibit a fibromyocyte-like transcriptional state before lipid loading, and that agLDL induces lipid-associated transcriptional programs in SMC-derived foam cells that continue to express an overall fibromyocyte gene expression pattern.

### Identification of Candidate Markers of SMC-Derived Foam Cells

To identify specific candidate markers of SMC-derived foam cells, we next focused on genes enriched in the putative single-cell RNA-seq–defined SMC foam cell cluster (Figures 1-3).

Candidate genes were prioritized based on differential expression criteria of high prevalence within the fibromyocyte cluster (pct.1 > 0.39) and low expression across other cell populations (pct.2 < 0.25) (Supplemental Table 2). Several genes, including *COL1A1*, *COL3A1*, *VCAN*, *ASPN*, and *OMD*, were not further pursued due to their extracellular matrix localization, which limits cell-specific interpretation in downstream validation. Additional candidates such as *PRSS23* and *CRLF1*, both encoding secreted proteins with less clearly defined roles in atherosclerosis, and *ADH1B*, a metabolic enzyme with broad expression, were also not prioritized. From this refined set, *CFH*, encoding complement factor H (CFH) and *SERPINE1*, encoding plasminogen activator inhibitor-1 (PAI-1), were selected as potential SMC foam cell markers (Figure 4&B). *CFH* showed concordant upregulation in agLDL-loaded SMCs (Supplemental Table 4) and has been reported in SMC-derived foam cell–associated gene sets,^18^ while *SERPINE1* has established roles in vascular remodeling and atherosclerosis.^39^

**Figure 4.**
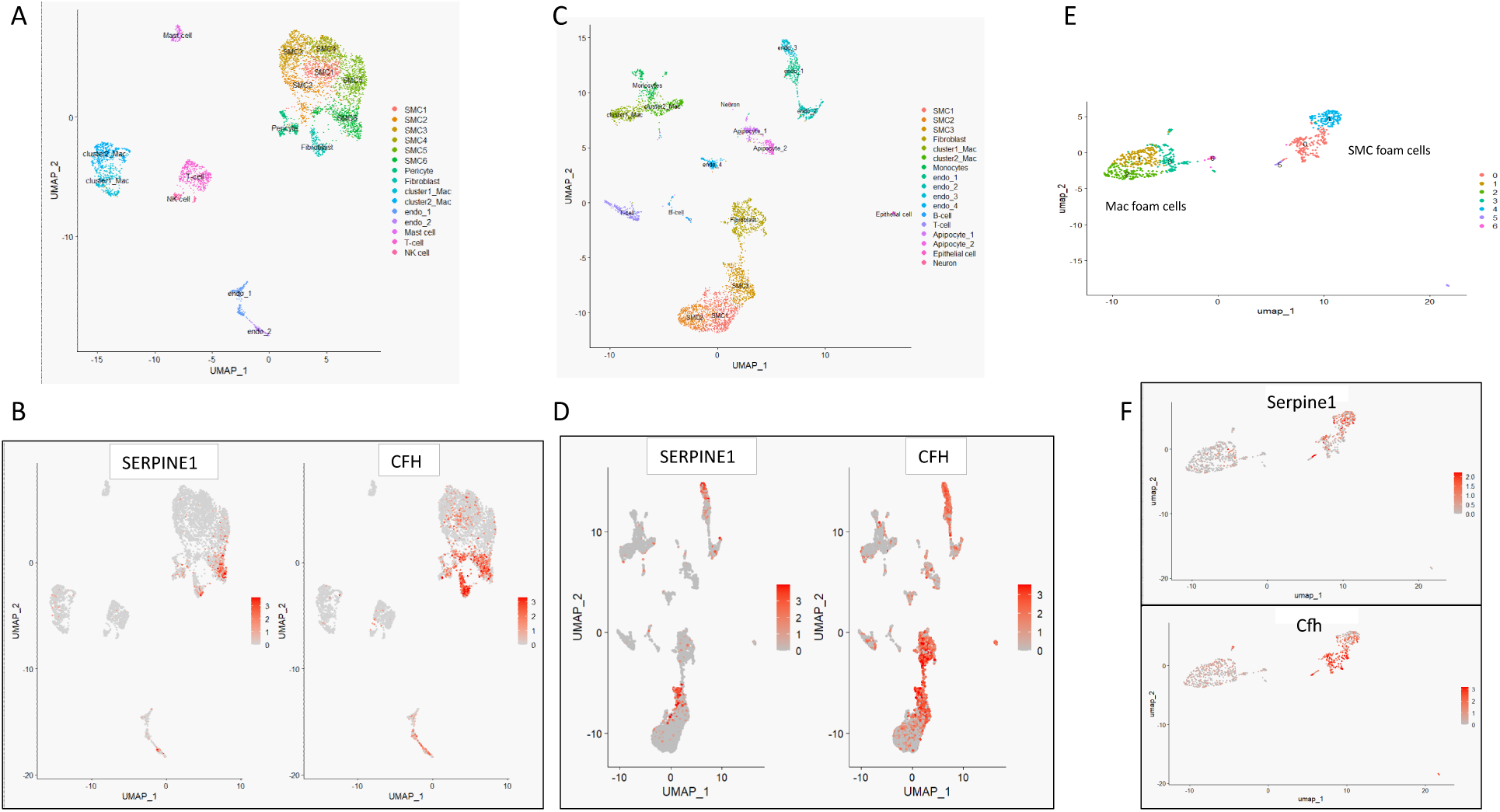
Identification and validation of SMC-derived foam cell markers across datasets. **(A)** UMAP projection of the integrated single-cell RNA-seq dataset showing annotated cell populations, including SMC subclusters, fibroblasts, endothelial cells, macrophages, and immune cells. **(B)** Feature plots displaying expression of candidate SMC foam cell– associated markers (*SERPINE1* and *CFH*) across all cell populations, demonstrating enrichment within fibromyocytes. **(C)** UMAP projection of an independent publicly available single-cell RNA-seq dataset from three human coronary artery samples, showing major vascular and immune cell populations annotated based on canonical marker expression. **(D)** Feature plots of *SERPINE1* and *CFH* expression in the independent dataset, confirming similar spatial enrichment patterns within the SMC compartment. **(E)** UMAP of foam cell populations derived from aortic cells isolated from atherosclerotic ApoE⁻/⁻ mice, highlighting distinct clusters of macrophage-derived foam cells and SMC-derived foam-like cells based on transcriptional profiles. **(F)** Feature plots of *Serpine1* and *Cfh* expression in the mouse dataset, demonstrating preferential enrichment in SMC-derived foam-like cells compared to macrophage foam cells.

To further validate these candidate markers, we next examined their expression in a publicly available human left coronary artery single-cell transcriptomic dataset from the Quertermous group (IGVF; dataset accession IGVFDS2114XGWE) (Figure 4C&D). One sample (P4) was excluded from this analysis because it contained heavily calcified lesions. Cell type annotation at level 1 was performed using canonical lineage markers and confirmed by DotPlot and FeaturePlot analyses (Supplemental Figure 4A-C). Subsequently, level 2 annotation of SMC subpopulations was carried out using the Mosquera atlas framework,^18^ enabling identification of contractile, fibromyocyte, and fibrochondrocyte SMC states (Supplemental Figure 4D). Within this framework, the SMC3 cluster was identified as the fibromyocyte population. Module scores generated using the literature-curated SMC foam cell gene set (Table 1) and genes upregulated in agLDL-loaded SMCs also showed strongest enrichment in the fibromyocyte cluster, with additional signal in fibroblasts (Supplemental Figure 4E, F). These patterns mirror the results shown in Figures 2A and 3C and further support fibromyocytes as encompassing the principal SMC population exhibiting foam cell transcriptional features. Consistent with this enrichment pattern, feature plots demonstrated that *SERPINE1* expression was enriched predominantly in the SMC3 (fibromyocyte) population. *CFH* expression was enriched in SMC3 (fibromyocytes) and was also detected in fibroblast populations, consistent with the pattern observed in our own dataset (Figure 4D). Importantly, neither *SERPINE1* nor *CFH* showed appreciable expression in macrophage clusters, supporting their association with non-macrophage foam cell–like states and reinforcing their candidacy as specific markers of SMC-derived foam cells.

To provide cross-species validation, we next analyzed a publicly available scRNA-seq dataset of aortic foam cells isolated from atherosclerotic ApoE⁻/⁻ mice ^2^ (GSM3215436) (Figure 4F). Unsupervised clustering of these cells revealed two major populations corresponding to Cd45⁺ immune-derived foam cells and Cd45⁻ non-immune foam cells, consistent with macrophage- and SMC-derived foam cell populations, respectively. Cell identity was further supported by canonical lineage markers (Supplemental Figure 5). Within this dataset, *Serpine1* expression was enriched in the Cd45⁻ SMC-derived foam cell population, with minimal expression in Cd45⁺ macrophage foam cells. Similarly, *CL* expression was preferentially detected in SMC-derived foam cells, with little to no expression in macrophage-derived foam cells. These findings recapitulate our observations in human datasets and support *SERPINE1* and *CFH* as conserved markers of SMC-derived foam cell states across species.

### Spatial Transcriptomic Validation of SMC Foam Cell Markers

To further validate these SMC foam cell candidate markers, we examined their spatial expression patterns in human coronary artery tissue using Xenium-based spatial transcriptomics. Fibromyocyte, macrophage, and macrophage foam cell module scores were first established based on previously defined gene signatures (Supplemental Figure 6A–C), enabling spatial delineation of SMC-derived and macrophage-associated transcriptional programs within the lesion. Consistent with these module scores, *CFH* and *SERPINE1* expression localized predominantly to regions enriched for fibromyocyte signatures, particularly within areas of high fibromyocyte score (Figure 6A; red box). In contrast, regions characterized by high macrophage foam cell module score (blue box) showed minimal expression of both *CFH* and *SERPINE1*, indicating that these markers are not associated with macrophage-derived foam cell populations. The presence of *CFH*-expressing fibromyocytes that lack *SERPINE1* expression (green box) indicates these two genes are regulated differentially within fibromyocytes. In parallel, THY1, previously suggested as a potential SMC foam cell marker,^19^ showed low level spatial enrichment in fibromyocyte-high regions.

Together, these spatial data, supported by the independently defined module scores in Supplemental Figure 6, demonstrate that *SERPINE1* and *CFH* expression, and to a small extent, *THY-1*, are specifically associated with fibromyocyte-rich regions and are spatially segregated from macrophage foam cell–enriched areas (Figure 5B). These findings provide spatial validation that SMC-derived fibromyocytes harbor distinct foam cell–associated transcriptional features that are molecularly and anatomically distinct from macrophage foam cells.

**Figure 5.**
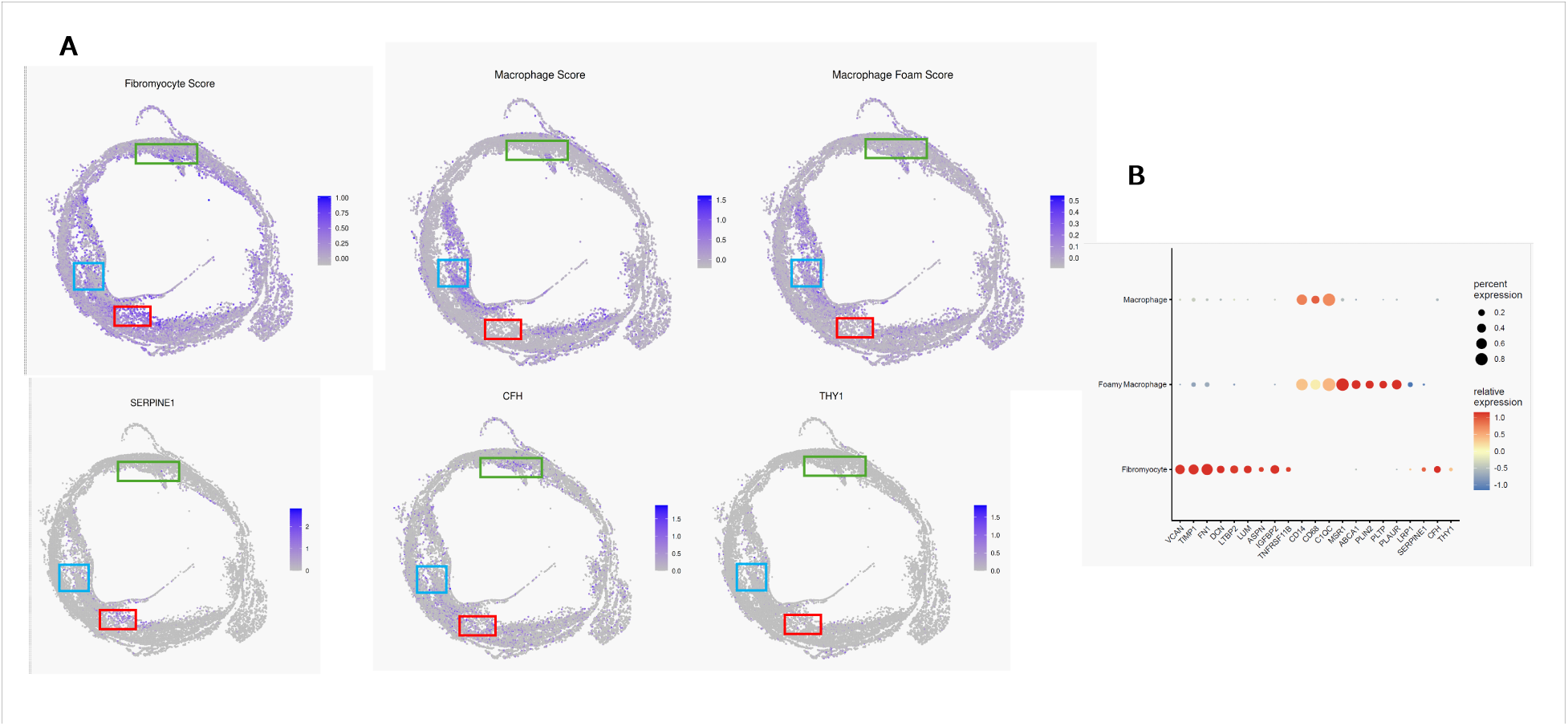
Spatial mapping of fibromyocyte and macrophage-associated programs in human coronary artery by Xenium. **(A)** Spatial feature plots showing module scores and gene expression overlaid on Xenium tissue coordinates. Fibromyocyte, macrophage, and macrophage foam cell gene signatures are visualized alongside expression of *SERPINE1*, *CFH*, and *THY1*. Color intensity represents normalized expression or module score. Distinct spatial regions are highlighted: fibromyocyte-enriched areas (red and green boxes) and macrophage-enriched areas (blue box) illustrating spatial segregation of SMC-derived and macrophage-associated programs. **(B)** Dot plot summarizing expression of selected marker genes across annotated cell populations in n=4 coronary segments from 4 separate explanted hearts. Dot size indicates the percentage of cells expressing each gene, and color represents scaled average expression.

**Figure 6.**
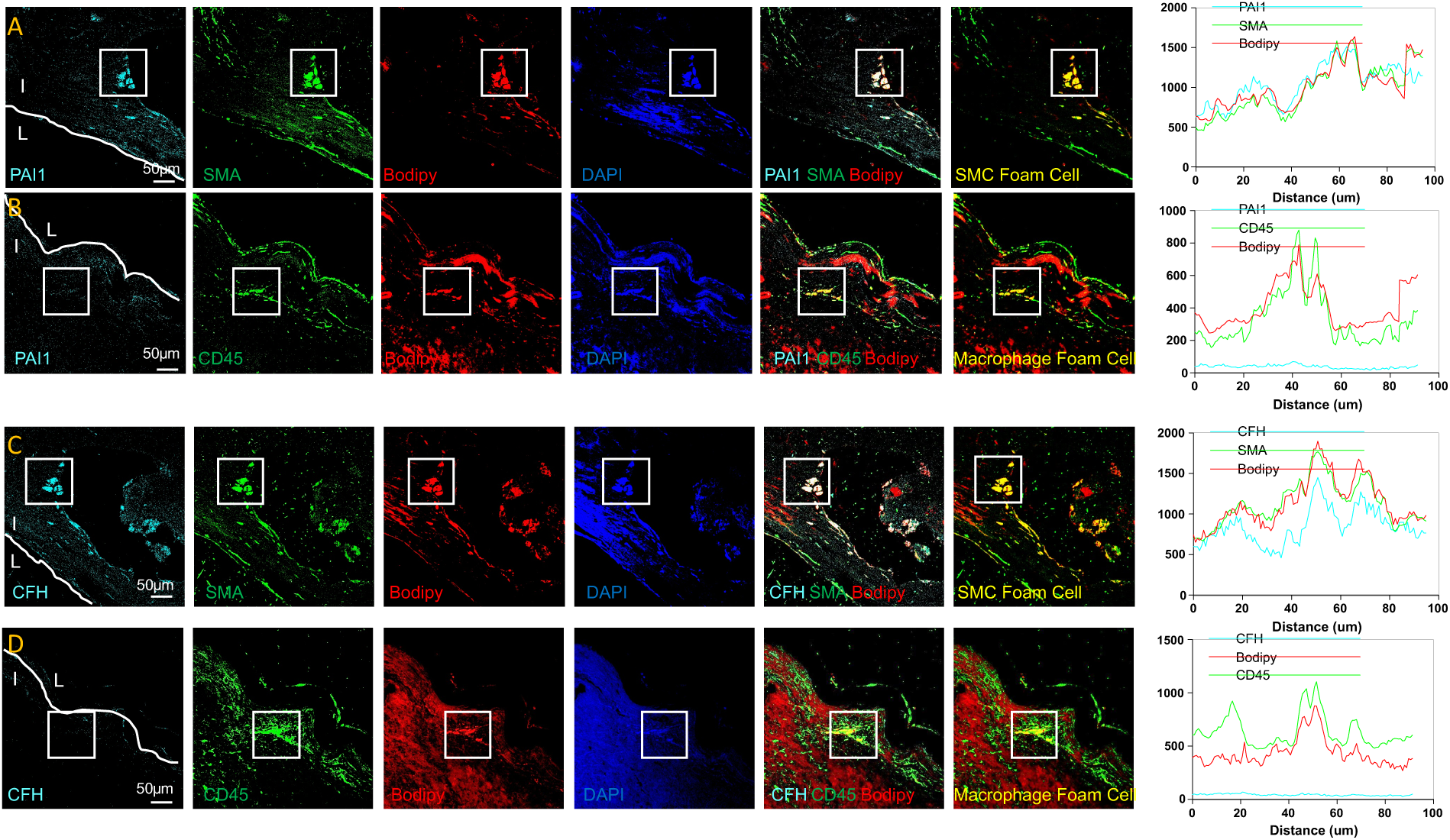
In situ validation of SERPINE1 (PAI-1) and CFH as candidate markers of SMC-derived foam cells in human coronary artery lesions. Serial sections of lipid-fixed human coronary artery plaques were stained for PAI-1 or CFH together with SMA, CD45, BODIPY, and DAPI. Panels **A** and **C** show representative regions in which PAI-1 or CFH co-localize with SMA and BODIPY, consistent with expression in SMC-derived foam cells. Panels **B** and **D** show regions stained with CD45 and BODIPY, together with PAI-1 or CFH, respectively, to assess macrophage foam cells. Merged images and highlighted overlays demonstrate co-localization patterns. Line-scan intensity plots across selected regions show overlap of PAI-1 or CFH with SMA and BODIPY, whereas little overlap is observed with CD45. I, intima; L, lumen. Scale bars = 50 μm. Representative of n=3 coronary sections from 3 separate explanted hearts.

### Tissue-level validation of *SERPINE1* and *CFH* expression in SMC-associated foam cell regions

To validate candidate markers of SMC-derived foam cells at the protein level, we performed immunofluorescence staining of human coronary artery sections for plasminogen activator inhibitor 1 (PAI-1, the gene product of *SERPINE1*) and CFH in combination with SMA, CD45, and BODIPY. This approach was designed to distinguish lipid-laden SMC-derived foam cells (BODIPY and SMA-positive) from macrophage-derived (BODIPY and CD45-positive) foam cells within atherosclerotic lesions. PAI-1 staining was detected in BODIPY-positive foam cell regions and showed clear overlap with SMA-positive cells, supporting expression in SMC-derived foam cells. In contrast, PAI-1 signal was minimal in CD45-positive BODIPY-positive regions, suggesting absence in macrophage foam cells. Similarly, CFH staining was observed in lipid-rich areas and co-localized with SMA-positive foam cells, whereas little CFH signal was detected in CD45-positive foam cell regions. Line-scan intensity analysis across selected regions of interest further supported co-distribution of PAI-1 or CFH with SMA and BODIPY, but not with CD45.

Together, these findings provide tissue-level validation that PAI-1 and CFH are associated with SMC-derived foam cells in human coronary artery lesions, and support the transcriptomic identification of these genes as candidate markers of SMC foam cells.

### Regulatory Pathways Associated with CFH and SERPINE1 Expression and Distinct Phenotypes of Smooth Muscle Cell and Macrophage Foam Cells

TGFβ signaling and extracellular matrix remodeling pathways are established regulators of *SERPINE1* expression,^40,41^ while inflammatory and interferon-associated pathways have been implicated in *CFH* induction.^42^ Because *CFH* is a key regulator of the complement cascade, complement-related genes are often co-expressed with *CFH*, reflecting a broader complement-associated transcriptional program.^43^ To investigate mechanisms that may contribute to *CFH* and *SERPINE1* expression in SMC-derived foam cells, we examined transcriptional programs associated with these genes in agLDL-loaded cultured SMCs. Bulk RNA-seq of agLDL-treated SMCs revealed coordinated induction of genes involved in TGFβ signaling, extracellular matrix remodeling, metabolic and stress responses, complement-associated programs, and inflammatory/interferon signaling (Figure 7A). To determine whether these candidate programs are recapitulated in vivo, we generated module scores using literature-curated gene sets representing putative regulatory programs associated with *SERPINE1* and *CFH* expression. Both signatures were enriched predominantly in early fibromyocytes and fibromyocytes, with minimal enrichment in contractile SMC states (Figure 7B), indicating that these transitional SMC states are enriched for transcriptional programs associated with induction of *SERPINE1* and *CFH*.

**Figure 7.**
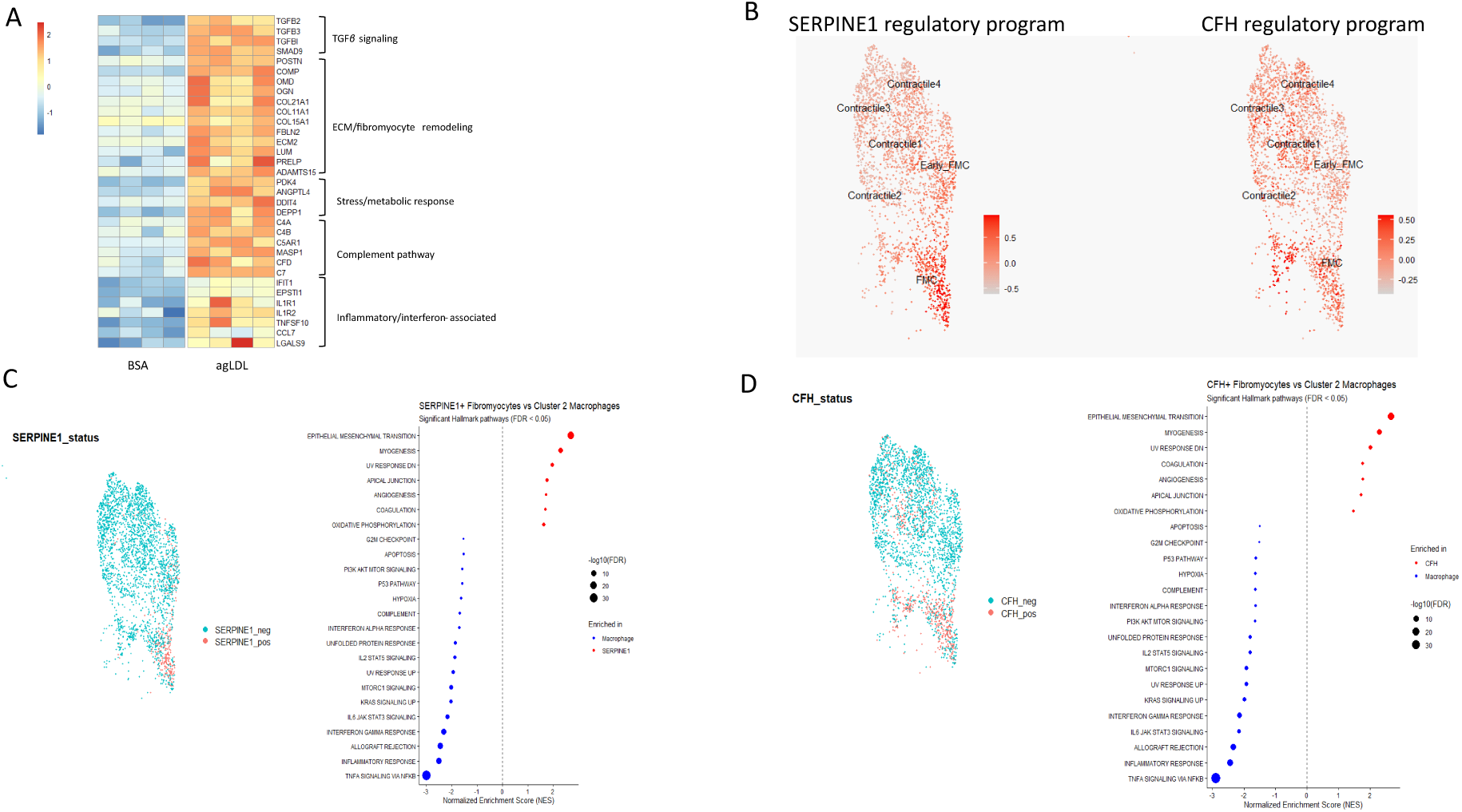
Putative regulatory pathways associated with SERPINE1 and CFH expression and divergent transcriptional phenotype of SMC-and macrophage-derived foam cells. **(A)** Heatmap of variance-stabilized bulk RNA-seq expression values from cultured human vascular SMCs treated for 72 hours with aggregated LDL (agLDL) or bovine serum albumin (BSA) control (n = 4 per group). Genes are grouped into functional categories representing candidate upstream pathways associated with *SERPINE1* (TGFβ signaling and ECM/fibromyocyte remodeling) and *CFH* (complement and inflammatory/interferon-associated pathways). Stress/metabolic response genes are included as broader cholesterol-induced programs that may contribute to regulation of both genes. Values are row-scaled (z-scores). **(B)** Feature plots showing module scores for curated *SERPINE1* and *CFH* regulatory gene programs across SMC subclusters in single-cell RNA-seq data. Both programs are enriched predominantly in Early fibromyocytes (Early_FMC) and fibromyocytes (FMC), with lower scores in contractile SMC states. **(C)** Left: UMAP highlighting *SERPINE1*-positive fibromyocytes. Right: Hallmark gene set enrichment analysis (GSEA) comparing *SERPINE1*-positive fibromyocytes with Cluster 2 macrophages. Positive normalized enrichment scores (NES; red) indicate pathways enriched in *SERPINE1*-positive fibromyocytes, whereas negative NES values (blue) indicate pathways enriched in macrophages. **(D)** Left: UMAP highlighting *CFH*-positive fibromyocytes. Right: Hallmark GSEA comparing *CFH*-positive fibromyocytes with Cluster 2 macrophages.

To define how SMC-derived foam cells differ from macrophage foam cells in vivo, we compared the top pathways activated in *CFH*-positive and *SERPINE1*-positive fibromyocytes with Cluster 2 macrophages, which showed the strongest enrichment for macrophage foam cell signatures. Gene set enrichment analysis demonstrated consistent enrichment of epithelial– mesenchymal transition, myogenesis, coagulation, angiogenesis, and oxidative phosphorylation pathways in *CFH*-positive and *SERPINE1*-positive fibromyocytes (Figure 7C–D). In contrast, Cluster 2 macrophages were enriched for inflammatory and immune-response pathways, including TNFα signaling via NF-κB, interferon responses, and IL6–JAK–STAT3 signaling.

Together, these findings suggest that cholesterol loading of SMCs activates transcriptional programs that are recapitulated in fibromyocytes in vivo and identify candidate regulatory and co-expression pathways associated with *SERPINE1* and *CFH* expression in SMC-derived foam cells. Strikingly, the top pathways activated in SMC foam cells are completely different from macrophage foam cells.

## DISCUSSION

In this study we utilized scRNA sequencing of coronary segments from patients with early to intermediate stage atherosclerosis to identify the distribution and phenotype of smooth muscle cell foam cells among the various clusters of coronary artery SMC subtypes, and validated these using additional coronary and carotid artery lesion datasets as well as bulk RNA sequencing of agLDL-loaded cultured SMCs. We further identified and validated specific markers of SMC foam cells using scRNA sequencing, spatial transcriptomics, and immunohistochemistry. Finally, we have begun to outline the biologic itinerary of SMC foam cells in relation to macrophage foam cells in lesions. Our findings indicate SMC foam cells in early to intermediate-stage atherosclerosis derive from fibromyocytes and have a unique gene expression pattern and biologic itinerary distinct from macrophage foam cells.

Using published markers of lipid-loaded cultured SMCs,^30,31^ we initially localized SMC foam cells to a cluster of modulated SMCs that show reduced expression of canonical markers of SMC differentiation and increased expression of extracellular matrix proteins, akin to fibroblasts, hence their designation fibromyocytes.^24^ Localization of SMC foam cells to this fibromyocyte cluster compared closely to that previously described by Mosquera *et al.* and Bashore *et al.* for “foam-like” or “foamy” SMCs.^18,19^ The accumulation of lipids within fibromyocytes is consistent with these modulated type of SMC populating the intima from early life and secreting the extracellular matrix proteins that bind apoB on atherogenic lipoproteins, providing a retained lipoprotein pool for uptake by fibromyocytes to form foam cells.^19^ Consistent with our in vitro findings, cultured human SMCs already exhibited a fibromyocyte-like transcriptional state before lipid loading, and agLDL induced the lipid-associated transcriptional program of SMC-derived foam cells that remains within the overall fibromyocyte transcriptional state. The localization of SMC foam cells or “lipomyocytes” to the SMC cluster 6/fibromyocytes as shown in Figures 1A and 1F combined with pseudotime trajectory analysis (Figure 1F) indicates further that these foam cells derive from more modulated SMCs along the trajectory of most to least differentiated SMCs (Figure 1F). In our analysis and those of Mosquera *et al.* and Bashore *et al.*,^18,19^ lipomyocytes showed a gene expression pattern very distinct from known markers of macrophages and macrophage foam cells (Figure 2). These comparisons suggest markedly different biologic pathways are activated in SMC and macrophage foam cells.

In vivo, agLDL is thought to be the most likely substrate for uptake by macrophages and SMCs to form foam cells.^36^ For many years, however, the method of loading SMCs with cholesterol bound to cyclodextrin has been adopted as a simpler and more efficient method to generate SMC foam cells than loading with lipoproteins in vitro.^33,34^ To further validate the distribution of SMC foam cells across SMC clusters, we examined whether loading of cultured SMCs with either agLDL or cholesterol bound to cyclodextrin (Chol-MβCD) induces gene expression programs that map to lipomyocytes observed in human coronary atherosclerotic tissue. As shown in Figure 3, agLDL induced a transcriptional profile that was clearly distinct from Chol-MβCD-treated SMCs (Figure 3A-3C). Importantly, the agLDL-induced gene signature mapped closely to the fibromyocyte/lipomyocyte region identified in the scRNA-seq atlas of coronary atheroma (Figure 2), supporting its relevance as an in vitro model of SMC-derived foam cell–like states (Figure 3C). In contrast, loading with cholesterol bound to cyclodextrin induced a markedly different transcriptional program that did not overlap at all with the lipomyocyte region but instead showed close similarity to macrophage-associated gene expression patterns. Consistent with this observation, gene ontology enrichment analysis revealed distinct biological processes associated with each treatment. agLDL preferentially induced pathways related to extracellular matrix organization, cell migration, and lipid-associated transport processes, whereas Chol-MβCD strongly enriched pathways linked to cytokine signaling, immune and inflammatory responses, and cell cycle/proliferation (Figure 3D). These results indicate that cholesterol loading via Chol-MβCD does not recapitulate the transcriptional state of SMC-derived foam cells observed in vivo. Similar findings have been reported by Conklin *et al*., showing in vitro loading of mouse vascular SMCs with Chol-MβCD did not mimic any SMC state found in mouse atheromas, although mouse SMC foam cells were not specifically examined.^35^ In contrast, loading with agLDL, or oxidized LDL,^30^ though somewhat less efficient than Chol-MβCD in inducing lipid accumulation in vitro, generates a gene expression program that closely reflects the lipomyocyte phenotype present in human coronary atheromas. This divergence reflects fundamental differences in the mode of cholesterol delivery by these two methods. Endosomal/lysosomal uptake of lipoproteins including agLDL results in the sequestration of lipoprotein-derived cholesteryl esters in the lysosomes of SMCs, due to relative deficiency of lysosomal acid lipase when compared to macrophages.^14^ Free cholesterol delivered to cells using Chol-MβCD binds directly the plasma membrane, bypasses lysosomes and can quickly be incorporated across the cell and induce endoplasmic reticulum stress, oxysterol generation, and induce an inflammatory response that closely resembles macrophage activation states. These results indicate a far less “macrophage-like” state of lipomyocytes than has been previously proposed, based on the gene expression profiles induced by Chol-_MβCD._10,33,34

To date, distinguishing lipomyocytes from macrophage foam cells in lesions has relied on SMC foam cells not expressing the pan-leukocyte marker CD45.^2,8,19^ In addition to lacking CD45 expression, having validated specific markers of lipomyocytes is critical to being able to distinguish key sites of cholesterol accumulation in plaques and to determine the effects of novel treatments and gene manipulations on plaque foam cell composition and depletion. In this report we identified *SERPINE-1*, encoding plasminogen activator inhibitor 1 (PAI-1), and complement factor H (*CFH*) as unique markers expressed at high levels in SMC cluster 6, at low levels by other SMC subtypes, and not expressed by macrophages. Both these markers localized to fibromyocytes in our scRNA-seq UMAP (Figure 4A and 4B) and to the proposed foam cell cluster in an independent cohort of 3 early to intermediate human coronary lesions (Supplemental Figure 3 and Figure 3C and 3D). In addition, these markers are enriched when mapped onto the scRNA-seq map of isolated CD45-foam cells from apoE-deficient mice fed a high fat diet, but expressed at very low levels in macrophage foam cells from these mice (Figure 4E and F), suggesting *SERPINE1* and *CFH* are conserved markers of lipomyocytes across species. To further validate these markers, we mapped them in human coronary lesions using Xenium spatial transcriptomics. A limitation of this method was the absence of lipid staining on an adjacent section because the samples were obtained from formalin-fixed, paraffin-embedded tissues, in which lipid content is removed during tissue processing. Despite this, by using a panel of either fibromyocyte or macrophage and macrophage foam cell gene markers (Supplemental Figure 5), we were able to map these markers to fibromyocytes and not to macrophages or macrophage foam cells (Figure 5). Of note from this analysis, we observed that some putative lipomyocytes express both *SERPINE1* and *CFH* while others express only one but not both of these markers. This implies there are alternate pathways regulating the expression of these two genes in lipomyocytes. We also determined the expression of another proposed SMC foam cell marker (*THY-1*, encoding the GPI-anchored protein CD90).^19^ While expression of *THY-1* was low relative to the other two markers, it is another potential gene that may be used when trying to identify lipomyocytes in lesions.

Further confirmation of the presence of PAI-1 and CFH in lipomyocytes but not in macrophage foam cells was determined using immunofluorescence staining of these proteins in human coronary lesions. These studies confirmed expression of these markers in lipomyocytes using co-staining of the marker antibodies with SMA and BODIPY, but low or no expression in SMC non-foam cells, macrophage foam cells, and macrophage non-foam cells (the latter two identified using CD45 +/- BODIPY staining) (Figure 6).

Lastly, a key aim of this study was to attempt to determine the biologic itinerary of lipomyocytes relative to macrophage foam cells in the plaque. Gene set enrichment analysis of the top pathways activated in lipomyocytes are completely distinct from macrophage foam cells, rather than being “macrophage-like” (Figure 7). The robust epithelial mesenchymal transition signal indicates lipomyocytes contribute strongly to the production of extracellular matrix (ECM) proteins and therefore to lipoprotein retention, fibrous-cap matrix remodeling, intimal expansion and either stabilization or fibrosis of the plaque, depending on the ECM quality.^44^ Enrichment of myogenesis pathways in lipomyocytes suggests these cells retain smooth muscle lineage programs despite lipid accumulation, consistent with a persistent fibromyocyte-like state and lack of transdifferentiation into macrophages. Pathway enrichment related to oxidative phosphorylation indicates lipomyocytes may be contributors to lipid processing, reactive oxygen species generation, inflammatory activation and possibly calcific/fibrotic remodeling.^45^ An increased coagulation signal suggests they may generate a locally thrombogenic plaque with increased vulnerability for occlusion after fibrous cap rupture or erosion.^46^ An increased signal for angiogenesis suggests lipomyocytes contribute to intraplaque neovascularization, strongly linked to plaque progression and vulnerability due to intraplaque hemorrhage, free cholesterol deposition from red blood cell membranes, and expansion of the necrotic core.^47^ In addition to these pathways, the observation that lipomyocytes retain most of their lipoprotein-derived cholesteryl esters in lysosomes rather than in the cytoplasm,^14^ and have low expression of the cholesterol exporter *ABCA1*,^8^ has implications for the ability of these cells to release their excess cholesterol to apoAI or HDL and to undergo regression in the plaque. Further studies are needed to determine how activation of these lipomyocyte pathways contributes to the progression and vulnerability of atheromas, and whether lipomyocytes represent a regression-resistant pool of cholesterol in the plaque that makes them a specific target for novel therapies to mobilize their excess cholesterol.

Limitations of this study include the potential alteration in intimal SMC gene expression following viral myocarditis and other causes of dilated cardiomyopathy that led to heart transplantation in the subjects whose coronary arteries were used for scRNA-seq. While this effect cannot be ruled out, the mapping of our scRNA-seq results onto the Mosquera *et al*.

MetaPlaq atlas generated mainly using carotid and coronary samples from patients with a primary diagnosis of atherosclerosis^18^ did not reveal clustering of our SMCs outside their SMC subtypes. This close alignment with SMCs in the MetaPlaq atlas validates the use of these samples derived from patients with non-ischemic cardiomyopathy. In addition, our coronary samples were noncalcified, as indicated by the absence of a separate cluster of chondromyocytes, and represent early to intermediate stage atherosclerosis. We cannot rule out whether lipomyocytes derived from more advanced lesions, which tend to have relatively fewer foam cells, would show the same gene expression patterns and pathway activation as those in early to intermediate lesions.

In conclusion, these findings provide the first detailed localization and analysis of SMC foam cells or lipomyocytes within scRNA-seq mapping of SMC clusters, identify *SERPINE1* and *CFH* as unique markers of lipomyocytes to distinguish them from macrophage foam cells in human atherosclerosis, and indicate unique pathways by which lipomyocytes may contribute to plaque progression and vulnerability. These findings, plus the known defects in the ability of SMC foam cells to mobilize excess cholesterol, indicate further studies of the role of lipomyocytes in the plaque will be critical to understanding their importance as a unique target to reduce ischemic vascular disease.

## Acknowledgments

The authors thank Amrit Samra for excellent technical support. S.A. and G.A.F. conceived the study, designed the experiments, analyzed the data and wrote the article. S.A. performed the majority of the experiments. Y.M., P.X., V.B., and T.C. performed experiments and contributed to experimental design. P.C. and T.Q. provided scRNAseq data for human samples and contributed to project design and data interpretation. P.A.H. and C.L.M. contributed to bioinformatic analysis and data interpretation. T.Q., D.Y.L., and M.W. provided human coronary tissues for Xenium analysis. U.T.A., M.T, M.K., and J.P.L. performed sample cutting, runs and quality control for Xenium analysis. A.B., T. Ö. and M.U.K. performed, analyzed, and provided data interpretation for Xenium analysis. We acknowledge the Single Cell Genomics Core at the University of Eastern Finland, supported by Biocenter Kuopio and Biocenter Finland, for providing spatial transcriptomics services. All authors have read and approved the article.

## Supplemental Material

Supplemental Methods

Supplemental Figures 1-6

Supplemental Data Table 1 Clinical Characteristics

Supplemental Data Table 2 All Markers

Supplemental Data Table 3 Gene Ontology SMC Clusters

Supplemental Data Table 4 DESeq2 agLDL vs Control

Supplemental Data Table 5 DESeq2 MBCD vs Control

Supplemental Data Table 6 agLDL up bulk gProfiler

Supplemental Data Table 7 CholMβCD up bulk gProfiler

## Nonstandard Abbreviations and Acronyms

BSA: fatty acid free bovine serum albumin
CE: cholesteryl ester
CFH: complement factor H
Chol-MβCD: cholesterol-methyl-β-cyclodextrin
FMC: fibromyocyte
GSEA: Gene Set Enrichment Analysis
IGVF: Impact of Genomic Variation on Function
PAI-1: plasminogen activator inhibitor 1
scRNA-seq: single cell RNA sequencing
SMC: smooth muscle cell

