## Supplemental Materials for "Transcriptomic, Specific Marker, and Pathway Analysis of Smooth Muscle Cell Foam Cells Relative to Macrophage Foam Cells in Human Atherosclerosis"

### Supplemental Methods

### Single-Cell RNA Sequencing Data Processing and Analysis

Single-cell capture and library preparation were performed at the Sequencing Core of the UBC Biomedical Research Centre. Single cells were encapsulated using the 10x Genomics Chromium platform, and libraries were prepared according to the manufacturer's instructions. Sequencing reads were processed using Cell Ranger and aligned to the human reference genome hg19 (GRCh37). Count matrices from individual samples were aggregated using the Cell Ranger aggr pipeline without subsampling normalization.

Downstream quality control and analysis were performed in R using Seurat version 4.3.0. Genes detected in fewer than three cells were excluded during Seurat object creation. Cells expressing fewer than 200 genes were excluded initially, followed by additional filtering to retain cells with 500–10,000 detected genes, 500–100,000 unique molecular identifiers (UMIs), and mitochondrial transcript content below 15%.

Gene expression values were normalized using the NormalizeData function with the default LogNormalize method, which scales counts to 10,000 transcripts per cell followed by natural log transformation. Highly variable genes were identified using FindVariableFeatures with the variance-stabilizing transformation method (selection.method = "vst") and 2,000 variable features.

Data were scaled using the ScaleData function, with regression of mitochondrial transcript percentage and total UMI counts (percent.mito and nCount_RNA) to minimize unwanted technical variation. Principal component analysis was performed on the highly variable genes, and the first 25 principal components were used for nearest-neighbor graph construction, clustering, and UMAP visualization. Cells were clustered using the shared nearest-neighbor modularity optimization algorithm implemented in FindClusters with a resolution of 0.6.

Cluster-specific marker genes were identified using FindAllMarkers, considering only positive markers expressed in at least 25% of cells (min.pct = 0.25) with a minimum log fold change of 0.25 (logfc.threshold = 0.25). Differential gene expression between selected clusters was performed using the MAST statistical framework, with p-values adjusted for multiple testing using the Bonferroni method.

**Cell Culture**

Human aortic smooth muscle cells (SMCs) were obtained from Gibco (catalog no. C-007-5C) and cultured in high-glucose Dulbecco's Modified Eagle Medium (DMEM; Cytiva HyClone) supplemented with Smooth Muscle Growth Supplement (Gibco catalog no. S00725). Cells were maintained under standard culture conditions at 37°C in a humidified incubator with 5% $CO_2$.

**Isolation and Aggregation of LDL**

LDL was isolated from the plasma of healthy human donors by gradient ultracentrifugation. LDL aggregates were prepared by vortexing in PBS for 60 seconds, as previously described.[5]

**Lipid Loading of Cells**

SMCs were cultured in DMEM supplemented with smooth muscle growth supplement until they reached 70–80% confluence. For aggregated LDL (agLDL) experiments, the medium was replaced with DMEM supplemented with 10% lipoprotein-deficient serum (LPDS) for 24 hours. Cells designated for cholesterol–methyl-β-cyclodextrin (Chol-MβCD) treatment were maintained in DMEM supplemented with smooth muscle growth supplement during this period. After two washes with PBS containing fatty acid-free bovine serum albumin (PBS-BSA), cells were incubated for 72 hours in DMEM containing 2 mg/mL BSA alone or with either 100 µg/mL aggregated LDL (agLDL) or 10 µg/mL Chol-MβCD.[6] Control cells were incubated for 72 hours in DMEM containing 2 mg/mL BSA alone.

**RNA Extraction**

Cells were harvested after 72 hours of treatment by washing once with PBS containing fatty acid-free albumin and twice with PBS, followed by detachment with Accutase. Cell pellets were collected by centrifugation, resuspended in a small volume of PBS, and freeze-dried prior to RNA extraction. Total RNA was then extracted using RNAzol RT according to the manufacturer's instructions and resuspended in 20 µL of DEPC-treated water for sequencing.

**Bulk RNA Sequencing Data Acquisition and Analysis**

RNA extracted from agLDL-loaded, Chol-MβCD-loaded, and non-loaded control smooth muscle cells was submitted to the Sequencing Core at the UBC Biomedical Research Centre for library preparation and sequencing. Bulk RNA-seq libraries were prepared using the Illumina Stranded mRNA Prep kit according to the manufacturer's instructions.

**RNA-seq Read Processing**

Sequencing reads were aligned to the Ensembl release 76 top-level human genome assembly using STAR (version 2.0.4b). Gene-level counts were generated from uniquely aligned, unambiguous reads using Subread (version 1.4.5). Transcript-level counts were generated using Sailfish (version 0.6.3). Sequencing quality was assessed using RSeQC (version 2.3), including total aligned reads, uniquely aligned reads, genes and transcripts detected, ribosomal RNA fraction, known junction saturation, and read distribution across annotated gene models.

**Differential Expression Analysis**

Differential expression analysis and principal component analysis were performed in R using DESeq2. Variance-stabilizing transformation was applied prior to principal component analysis, which was performed using the 500 most variable genes. Differential expression between loaded and non-loaded smooth muscle cells was conducted according to standard DESeq2 workflows. P-values were adjusted for multiple testing, and genes with an adjusted p-value < 0.05 were considered significantly differentially expressed.

**Publicly Available Human Coronary Artery Transcriptomic Datasets**

Publicly available transcriptomic datasets from human left coronary artery samples were obtained from the (IGVF) data portal (dataset accession: IGVFDS2114XGWE). Four samples were included in the analyses. Processed Seurat objects and associated metadata provided by the consortium were used for downstream analyses, including cell clustering, annotation, and comparative transcriptional profiling.

**Lipid Fixation**

Formalin-fixed human coronary artery tissues were processed to carefully preserve lipid content in paraffin sections as previously described.[10] In brief, formalin-fixed tissues (1–2 mm thick) were kept in an emulsion of linoleic acid (Fisher) and lecithin (Jamieson Natural Sources) in 70% ethylene glycol (Sigma) at 56°C for 3 to 5 days. Tissues were rinsed for at least 8 hours in several changes of 70% ethanol followed by several changes of distilled water, immersed in 2% chromic acid for 24 hours at 4°C, then rinsed in several changes of distilled water for 24 hours. Finally, tissues were placed in a 5% sodium bicarbonate solution for 24 hours followed by rinsing in water for at least 8 hours. Tissues were then embedded and cut in paraffin sections and stained with oil red O. This method preserves neutral lipids in tissues for subsequent lipid staining.

**Immunofluorescence and Confocal Microscopy**

Serial 7-µm sections of human coronary artery were prepared for immunofluorescent staining. Sections were blocked with 10% goat serum and subsequently incubated with primary antibodies against α–smooth muscle actin (SMA; Abcam, ab7817), plasminogen activator inhibitor-1 (PAI-1; Abcam, ab222754), complement factor H (CFH; Proteintech, 12748-1-AP), Fibulin 1 (Abcam, ab211536), or CD45 (R&D Systems, MAB1430). For detection, goat anti-rabbit Alexa Fluor 647 secondary antibody was used for PAI-1 and CFH, goat anti-mouse Alexa Fluor 555 secondary antibody was used for CD45 and SMA, and goat anti-mouse Alexa Fluor 647 secondary antibody was used for Fibulin 1. Foam cells were stained with 10 µg/mL BODIPY 493/503 (Invitrogen) concurrently with secondary antibody incubation. Fluorescence images were acquired using a Zeiss laser scanning confocal microscope and analyzed with ImageJ software.

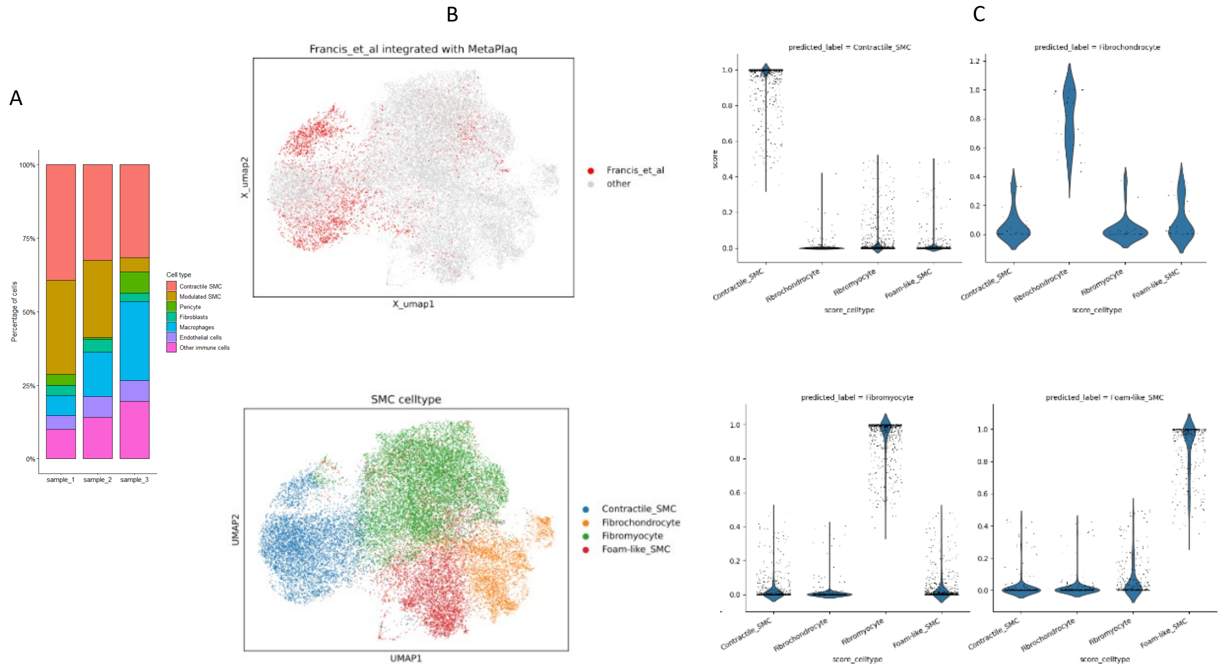


**Supplemental Figure 1. Cross-sample cell-type composition and validation of SMC state classification following integration with MetaPlaque.**

**(A)** Stacked bar plots showing the percentage of major cell types across individual samples (sample_1–sample_3). Cell identities were grouped into contractile SMCs, modulated SMCs, fibroblasts, macrophages, endothelial cells, and other immune cells, illustrating overall cellular composition and consistency across samples. **(B)** Integration of VSMC subpopulation with MetaPlaq v2 using the scArches approach (Unpublished). VSMCs are colored in red in the top panel and MetaPlaq v2 cells are in gray in low-dimensional space. Low-dimensional embeddings are colored by VSMC subtype in the lower panel. **(C)** SCANVI predicted probability of label transfer in VSMC subtypes for individual cells. The SCNAVI predicted annotation is shown on the x-axis and is plotted against the probability of annotation on the y-axis. As shown, predicted annotations show high probabilities for their corresponding label and other subtypes show low probability scores for other annotations.

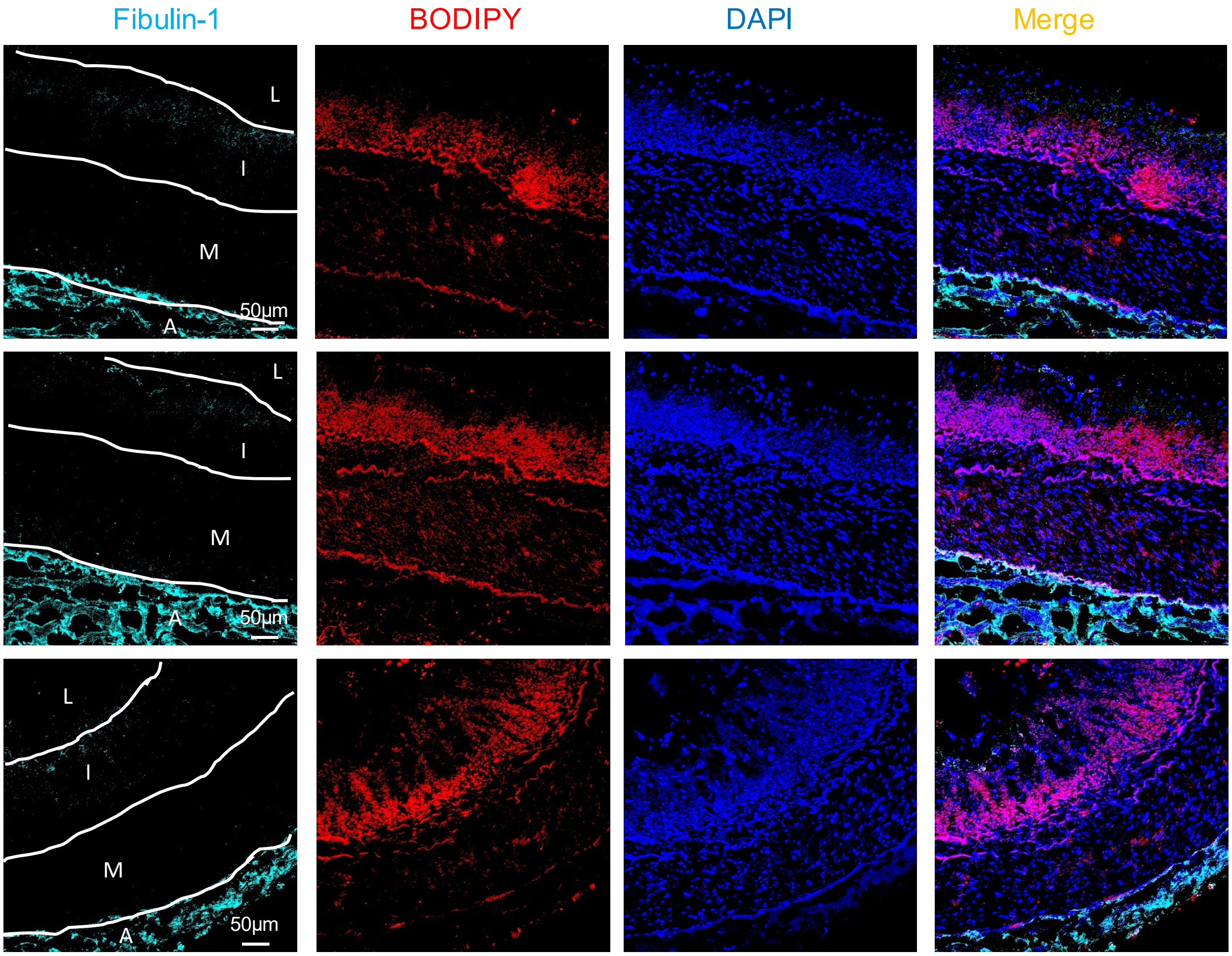


**Supplemental Figure 2. Lack of lipid accumulation in Fibulin-1–positive fibroblasts in human coronary artery.**

Representative immunofluorescence images of human coronary artery sections stained for Fibulin-1 (cyan) to identify fibroblasts, neutral lipid using BODIPY 493/503 (red), and nuclei with DAPI (blue). Merged images are shown in the right column. Across multiple regions, Fibulin-1–positive fibroblasts are localized primarily within the adventitia and do not exhibit co-localization with BODIPY-positive lipid droplets, which are predominantly observed in the intimal lesion. Tissue layers are indicated as lumen (L), intima (I), media (M), and adventitia (A). Scale bars, 50 µm.

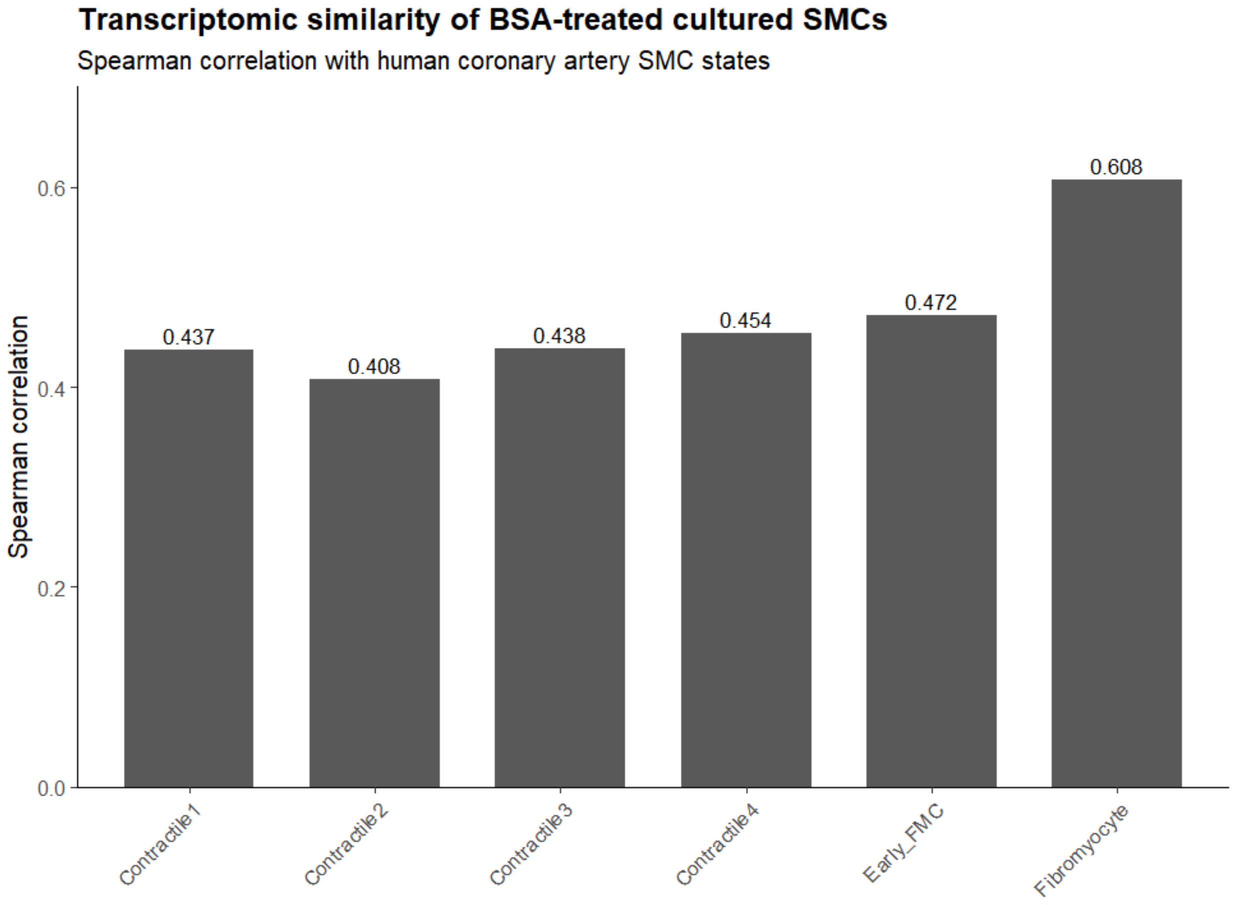


**Supplemental Figure 3. Transcriptomic similarity of BSA-treated cultured SMCs to human coronary artery SMC states.**

Bulk RNA-seq was performed on BSA-treated cultured human vascular SMCs. The average transcriptomic profile of BSA-treated SMCs was compared with pseudobulk gene expression profiles generated from each human coronary artery SMC cluster identified by single-cell RNA sequencing. Spearman correlation coefficients were calculated using the top 2,000 highly variable genes shared between the bulk RNA-seq and pseudobulk datasets. Bars represent the Spearman correlation coefficient between the BSA-treated bulk RNA-seq profile and each SMC cluster pseudobulk profile.

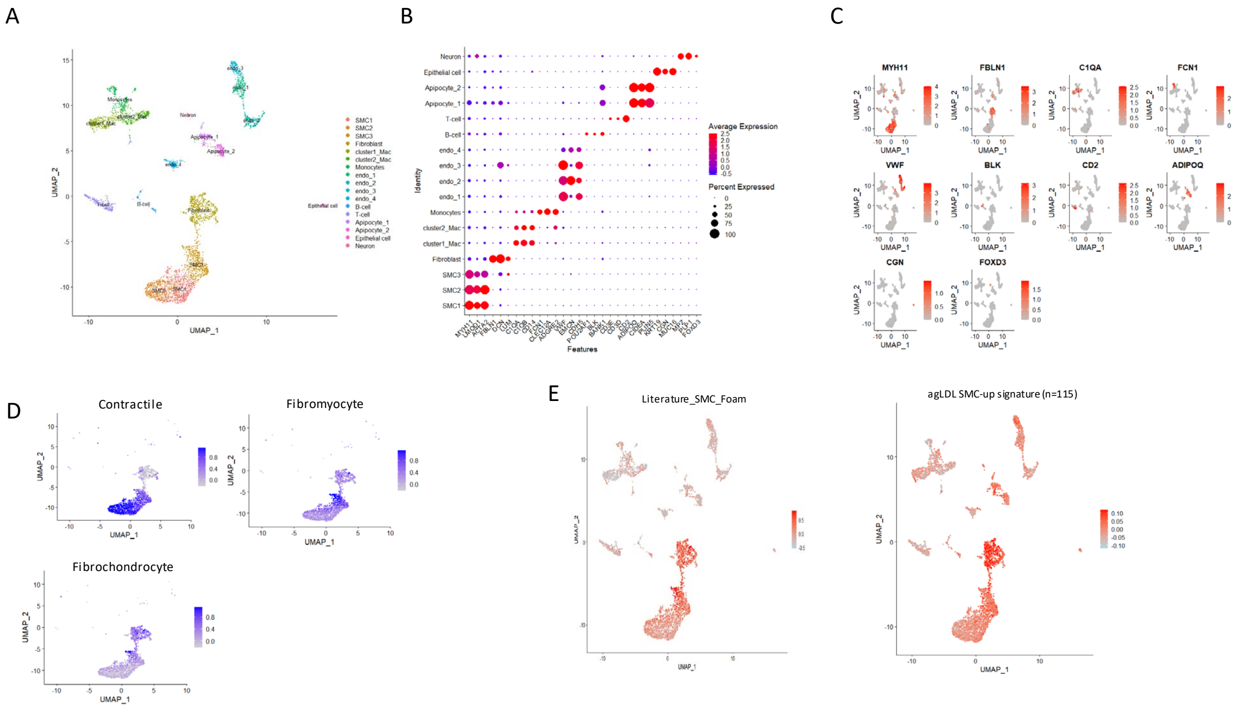


**Supplemental Figure 4. Validation of vascular cell type annotation and SMC state programs in an independent public dataset.**

**(A)** UMAP projection of a publicly available single-cell RNA-seq dataset generated from three human coronary artery samples, showing major vascular and immune cell populations identified by unsupervised clustering and annotated based on canonical marker expression.

**(B)** Dot plot of canonical lineage markers used for Level-1 cell type annotation in the validation dataset. Dot size represents the percentage of cells expressing each gene, and color indicates average scaled expression. **(C)** Feature plots of representative marker genes confirming the identity of major cell populations, including smooth muscle cells (*MYH11*), endothelial cells (*VWF*), immune cells (*C1QA*, *CD2*), fibroblasts (*FBLN1*), and other lineage-specific markers.

**(D)** UMAP projections displaying module scores for previously defined SMC transcriptional programs (contractile, fibromyocyte, and fibrochondrocyte), demonstrating conservation of SMC phenotypic states in the independent dataset. **(E)** AddModuleScore analysis using a literature-curated SMC foam cell–associated gene set, showing enrichment of the SMC foam-like program across the UMAP. **(F)** AddModuleScore analysis using genes upregulated in agLDL-loaded SMCs from bulk RNA-seq (n = 115 genes), showing projection of the agLDL-induced SMC signature onto the single-cell UMAP.

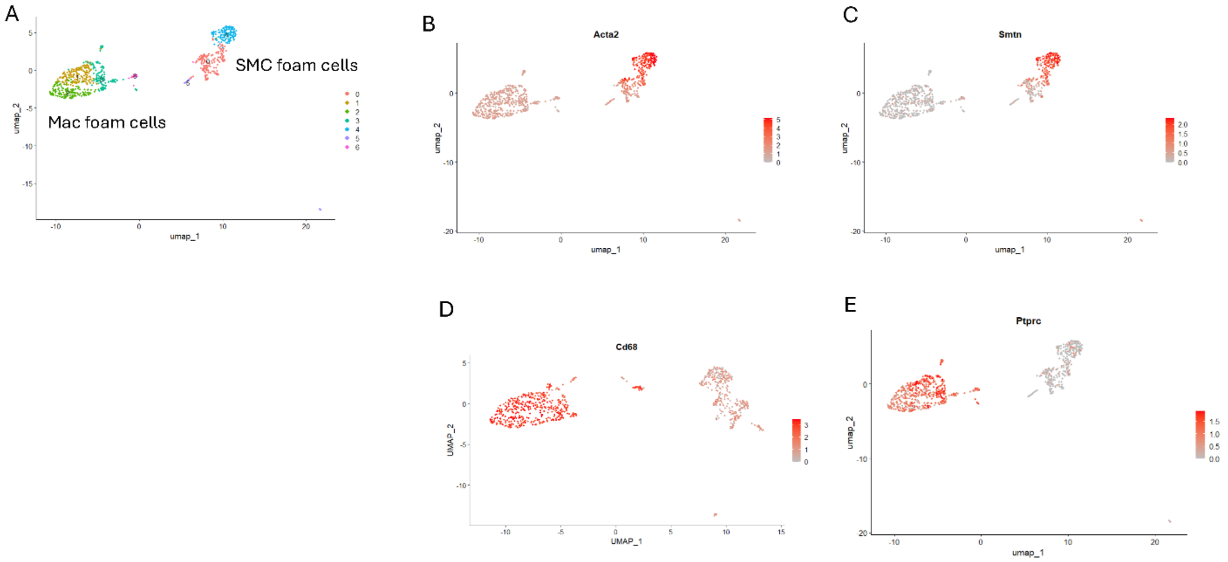


**Supplemental Figure 5. Identification of SMC-derived and macrophage-derived foam cells in a mouse aortic single-cell RNA-seq dataset.**

(**A)** UMAP visualization of single-cell RNA-seq data from BODIPY$^{hi}$ foam cells isolated from atherosclerotic ApoE knockout mouse aortas (GSM3215436), revealing two major populations corresponding to macrophage-derived foam cells and non-immune foam cells consistent with SMC-derived foam cells. **B–C**, Feature plots showing expression of the canonical smooth muscle markers *Acta2* and *Smtn*, confirming enrichment within the SMC-derived foam cell population. **(D–E)** Feature plots showing expression of the macrophage markers *Cd68* and *Ptprc* (Cd45), which are enriched in the macrophage-derived foam cell population.

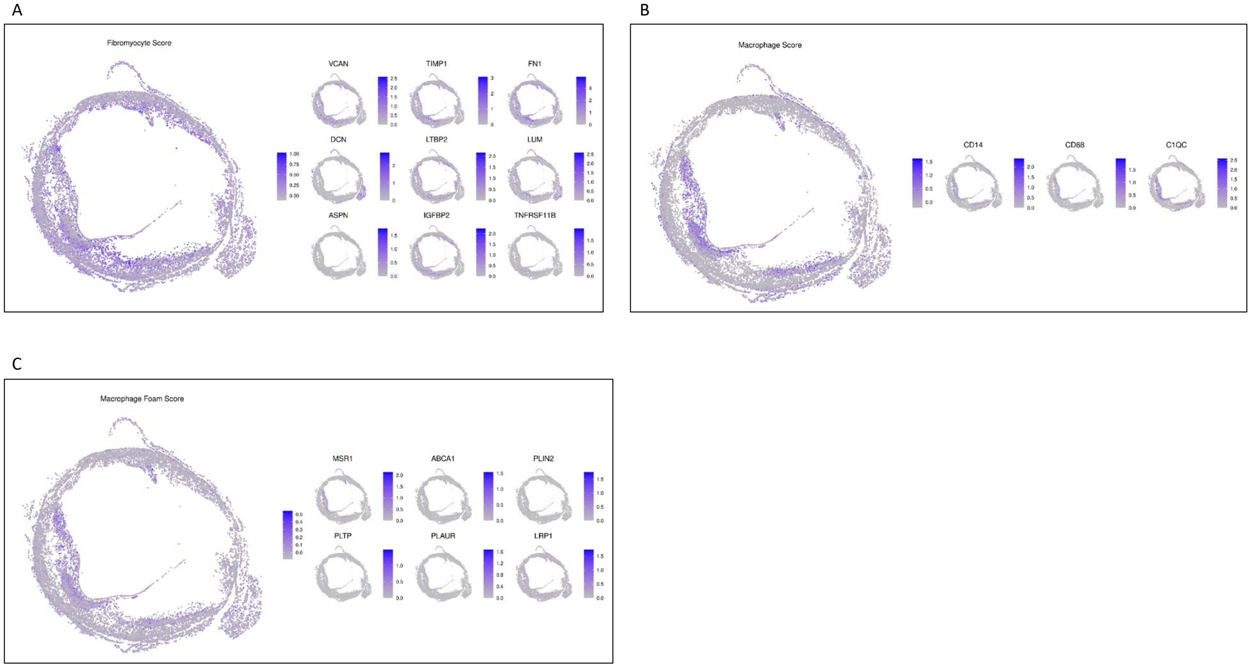


**Supplemental Figure 6. Spatial mapping of fibromyocyte-, macrophage-, and macrophage foam cell-associated gene programs in human coronary artery by Xenium.**

**(A)** Spatial distribution of the fibromyocyte module score, calculated using extracellular matrix and remodeling-associated genes (*VCAN, TIMP1, FN1, DCN, LTBP2, LUM, ASPN, IGFBP2, TNFSF11B*), highlighting regions enriched for fibromyocyte-like smooth muscle cell states. **(B)** Spatial distribution of the macrophage module score, calculated using canonical macrophage markers (*CD14, CD68, C1QC*), demonstrating localization of macrophage populations within the lesion. **(C)** Spatial distribution of the macrophage foam cell module score, calculated using lipid handling and foam-associated genes (*MSR1, ABCA1, PLIN2, PLTP, PLAUR, LRP1*), indicating enrichment of lipid-loaded macrophages in macrophage-dense regions. Color intensity represents normalized expression or module score. These data demonstrate spatially distinct fibromyocyte and macrophage-associated transcriptional programs within the atherosclerotic plaque.
